# Gut and nasal microbiome trajectories shape respiratory disease severity in calves exposed to antibiotic and pathogen challenge

**DOI:** 10.64898/2026.10.16.759607

**Authors:** Maverick Monie--Ibanes, Thomas Peyret, Güénaelle Derolez, Marie Corbin, Florence Mompart, David Bars-Cortina, Fatima-Zohra Sikht, Christine Citti, Xavier Nouvel, Eric Baranowski, Gilles Meyer, Mariette Ducatez, Núria Mach

## Abstract

Bovine respiratory disease complex (BRDC) is a multifactorial syndrome resulting from interactions among pathogens, host responses, and environmental stressors. However, substantial variation in disease severity persists among animals exposed to similar infectious challenges. To assess the contribution of the respiratory and gut microbiota to BRDC severity, we experimentally inoculated 28 calves with *Mycoplasma bovis*, bovine coronavirus (BCoV), and influenza D virus (IDV), with half receiving metaphylactic tilmicosin. Clinical trajectories were classified into mild, moderate, and severe phenotypes using unsupervised clustering of longitudinal health scores.

Severe BRDC was associated with higher and more persistent nasal IDV loads, yet pathogen burden alone failed to explain the marked inter-individual variability in clinical outcomes. Tilmicosin-treated calves developed substantially milder disease despite comparable pathogen loads, coinciding with lower systemic inflammation, suggesting that host-pathogen interactions, rather than pathogen replication alone, influenced disease severity.

Neither nasal nor faecal microbiota diversity consistently explained BRDC severity. However, calves that later developed severe disease displayed a more homogeneous nasal microbiota before infection, indicating that early microbial community structure may influence susceptibility. In contrast, the strongest disease-associated microbial signatures were observed in the faecal microbiota. Gut antibiotic-induced dysbiosis revealed associations between BRDC severity, specific intestinal bacterial taxa, systemic inflammation, and faecal short-chain fatty acid (SCFA) profiles. Notably, *Chlamydia* was strongly associated with disease severity, while several other taxa were linked to inflammatory biomarkers and SCFA concentrations.

Collectively, these findings support a holobiont-driven model of BRDC in which disease severity emerges from interactions among pathogens, host inflammatory responses, and microbiota-associated functions. The stronger associations observed in the intestinal microbiota, along with links between gut bacteria, SCFAs, and disease outcomes, provide evidence for a functional gut-lung axis in calves and highlight microbial metabolism as a potential target for strategies to improve respiratory health and reduce antibiotic use.

**Implications:** Bovine respiratory disease is one of the most costly health and welfare problems in cattle production. This study shows that disease severity is not determined only by pathogen infection but is also linked to changes in the gut microbiome and its metabolic activity. Calves with different microbiome profiles developed markedly different clinical outcomes despite similar pathogen exposure. The findings provide new evidence for a gut-lung connection in cattle and identify microbial functions associated with disease resilience. These results may support the development of microbiome-based strategies to improve animal health, reduce production losses, and decrease reliance on antibiotics in livestock systems.

## Introduction

Bovine Respiratory Disease Complex (BRDC) is one of the most significant health and welfare challenges affecting cattle production worldwide [1]. The disease is associated with substantial economic losses resulting from mortality, reduced growth performance, treatment costs, and long-term effects on productivity [2, 3]. BRDC accounts for approximately 14-20% of pre-weaned calf mortality and can contribute to even higher losses in specific production systems [2, 3]. Despite decades of research, BRDC continues to represent a major challenge owing to its multifactorial nature and the complex interactions occurring among pathogens, host responses, environmental conditions, and management practices [1, 4–8]. Transportation, commingling, dietary changes, and weaning induce physiological stress and immune dysregulation, increasing susceptibility to respiratory pathogens. Because outbreaks frequently occur following transportation to feedlots, BRDC is often referred to as "shipping fever"[9].

BRDC is generally initiated by viral pathogens including Bovine Respiratory Syncytial Virus (BRSV), Bovine Coronavirus (BCoV), Influenza D Virus (IDV), and Bovine Parainfluenza Virus type 3 (BPIV-3), which predispose the respiratory tract to secondary bacterial colonisation [1, 10–12]. The bacterial species most frequently associated with disease include *Pasteurella multocida, Mannheimia haemolytica, Histophilus somni*, and *Mycoplasma bovis* [5, 13]. Because multiple pathogens frequently coexist within the respiratory tract, BRDC is widely recognised as a polymicrobial disease, with mixed viral-bacterial infections often associated with the most severe clinical outcomes [1, 14].

Although pathogen exposure is a prerequisite for disease development, a major unresolved question in BRDC research is why animals exposed to similar infectious and environmental pressures often develop markedly different clinical outcomes. Under field and experimental conditions, some calves remain clinically healthy or show only mild respiratory signs, whereas others develop severe, prolonged disease [15–17]. Several studies have linked pathogen burden to disease severity, suggesting that higher pathogen replication may contribute to worse outcomes [16, 18]. However, these relationships are often inconsistent, and calves harbouring similar pathogen loads can exhibit strikingly different clinical trajectories. Such observations indicate that pathogen presence and abundance alone cannot fully explain disease severity and suggest that host-associated factors play a critical role in determining resilience or susceptibility to BRDC [19].

Increasing evidence indicates that microbial communities inhabiting mucosal surfaces are important regulators of these host responses, influencing both immune homeostasis and resistance to pathogen colonisation [20–23]. Among these communities, the respiratory microbiota has emerged as a potentially important determinant of respiratory health [24, 25]. The nasal microbiota is the first microbial ecosystem inhaled pathogens encounter and plays a critical role in colonisation resistance, immune modulation, and maintenance of mucosal homeostasis [26]. Alterations in respiratory microbial communities are associated with respiratory disease in calves, suggesting that microbiota composition may influence pathogen establishment, host susceptibility, disease progression, and clinical outcome [18, 27–30].

Recent studies have further expanded our understanding of the development and dynamics of the calf respiratory microbiota and its response to environmental factors, management practices, antimicrobial treatments, and disease challenges [14, 15, 17, 18, 28, 31–39]. However, whether respiratory microbial communities actively influence the severity of BRDC after pathogen exposure remains largely unknown. Identifying microbial signatures associated with resistance or susceptibility to severe disease could provide novel opportunities for disease prediction and microbiome-based prevention strategies.

However, respiratory health is unlikely to be determined solely by airway microbial communities. Increasing evidence supports the existence of a bidirectional gut-lung axis, whereby intestinal and respiratory microbiotas interact through immune, metabolic, and microbial signalling pathways [40–42]. Through the production of microbial metabolites, particularly short-chain fatty acids, and the modulation of systemic immune responses, the gut microbiota can influence respiratory mucosal immunity, susceptibility to pathogen colonisation, and the outcome of respiratory infections [43–45]. Conversely, respiratory disease and inflammation may alter gut microbial composition, highlighting the interconnected nature of these mucosal ecosystems [41]. Although investigations in cattle remain relatively limited, emerging studies suggest that interactions between intestinal and respiratory microbial communities contribute to calf health, growth performance, and disease susceptibility, and may therefore represent an important but largely unexplored component of BRDC pathophysiology [9, 15, 17, 46]. Consequently, determining how respiratory and intestinal microbiotas jointly influence host responses to BRDC pathogens may provide novel insights into mechanisms of disease resistance and identify new targets for microbiome-based prevention strategies.

Antimicrobial treatment represents another major factor influencing BRDC outcomes [47]. While therapeutic and metaphylactic antimicrobials are widely used to reduce the morbidity and mortality associated with BRDC, their effects may extend beyond direct antibacterial activity [30, 33, 35, 48, 49]. Consequently, antimicrobial administration can shape disease outcomes through its combined effects on pathogens, host immunity, and the microbiota [50, 51]. Elucidating these interconnected mechanisms is particularly relevant in the context of efforts to develop microbiome-informed strategies for disease control and to reduce reliance on antimicrobials in cattle production.

Despite growing interest in the role of microbiota-host-pathogen interactions in BRDC, most studies have examined only specific components of this complex system. Previous studies have typically focused on either respiratory microbiota dynamics during BRDC [34, 52, 53], the coordinated development of respiratory and intestinal microbiotas in healthy calves [9, 15], or microbiota-based interventions aimed at reducing pathogen colonisation and improving respiratory health [28, 35, 36, 49, 54]. As a result, the mechanisms through which respiratory pathogens, host responses, and interconnected microbial ecosystems jointly contribute to disease resilience or susceptibility remain poorly understood.

In the present study, we combined experimental pathogen inoculation, longitudinal clinical assessment, pathogen quantification, and microbiota profiling to investigate the determinants of BRDC severity in calves. We hypothesised that administration of tilmicosin to healthy animals would induce respiratory and intestinal dysbiosis, thereby disrupting microbiota-mediated colonization resistance and host-microbe interactions. Consequently, we expected that calves exposed to similar pathogen challenges would exhibit substantial heterogeneity in clinical outcomes, with antibiotic-induced alterations of the microbiota contributing to differences in pathogen establishment, replication, and disease severity. We further hypothesised that pathogen burden alone would not fully explain this variability and that specific features of the nasal and intestinal microbial communities would be associated with either susceptibility or resilience to BRDC. By integrating pathogen dynamics, clinical trajectories, and respiratory and faecal microbiota profiles, we aimed to determine whether antimicrobial-induced microbiota disruption influences disease outcome and to identify microbial and biological factors associated with BRDC severity.

## Material and method

### Ethics approval

The local animal care and use committee and the French Minister of Research reviewed and approved the study protocol for the calves’ study (reference: APAFIS #49942-2024062000479999 v3, dated September 23rd, 2024, Ethics Committee no. 115). All protocols were conducted in accordance with the EEC Regulation (No. 2010/63/UE) governing the care and use of laboratory animals, which has been in effect in France since January 1, 2013.

### Experimental design

The present study was conducted using samples and metadata generated during the experimental calf coinfection trial described in detail by Méndez et al. (2026)[55]. Briefly, twenty-eight dairy calves (Montbéliard and Holstein) were reared under standardised conditions from birth at the INRAE Herbipole experimental farm (France) and subsequently transferred to the National Veterinary School of Toulouse (ENVT, France) for experimental procedures. Méndez et al. (2026) [47] provide detailed descriptions of calf management, feeding strategy, vaccination protocol, housing conditions, biosecurity measures, group allocation, animal welfare procedures, and ethical approvals are provided in Méndez et al. (2026) [55]. At ENVT, calves were allocated to four experimental groups according to infection and antimicrobial exposure status: (i) experimentally inoculated and antibiotic-treated, (ii) experimentally inoculated and non-treated, (iii) non-inoculated and antibiotic-treated, and (iv) non-inoculated and non-treated.

Antibiotic-treated calves received tilmicosin prior to pathogen challenge (12.5 mg/kg, twice daily; Tilmovet 250 mg/mL, Huvepharma, Antwerp, Belgium) from days -5 to -3 of the experiment. Antibiotic treatment was administered exclusively to the *nonInoc_ATB* and *Inoc_ATB* groups. Tilmicosin was administered to healthy calves as a microbiota perturbation strategy rather than as a therapeutic intervention. The experimental design was based on the hypothesis that antimicrobial-induced disruption of resident microbial communities could alter the respiratory and intestinal mucosal environment, thereby influencing susceptibility to subsequent pathogen colonization and infection. This approach was also intended to mimic commercial production practices, where metaphylactic antimicrobial treatments are frequently administered to clinically healthy calves during periods of elevated disease risk, such as weaning, transportation, commingling, or feedlot arrival [56, 57]. Calves received a single dose of tilmicosin 72 h before pathogen exposure, a delay expected to substantially reduce residual antimicrobial activity at the time of infection while maintaining antibiotic-induced alterations of the microbial ecosystem [58]. This model allowed us to evaluate whether prior antimicrobial exposure and the resulting dysbiosis could affect pathogen establishment, disease progression, and host-microbiota interactions following experimental challenge.

On day 0, calves assigned to the infected groups were experimentally coinoculated with three BRD-associated pathogens: IDV (strain D/bovine/France/5920/2014), BCoV (strain ENVT-66), and *M. bovis* (strain RM16), following the procedures described by Méndez et al. (2026) [55]. Briefly, viral strains were propagated in cell culture and titrated prior to challenge, whereas *M. bovis* RM16 was cultured in SP4 medium and quantified by colony-forming unit (CFU) counting. Inocula were prepared immediately before administration to ensure viability and consistency of the infectious dose.

Animals were housed in separate pens providing approximately 10.5 m² per calf. To prevent pathogen transmission, infected and non-infected groups were physically segregated by a polyethylene barrier and managed using strict biosecurity procedures, including dedicated personnel, equipment, and husbandry practices. Water was available *ad libitum*, and bedding was regularly renewed throughout the experiment.

### Feeding management

Throughout the study, calves were fed exclusively with a commercial milk replacer identical to that used before transportation to the ENVT. The amount supplied was adjusted weekly according to body weight to target an average daily gain of 0.8 kg/day. Milk was offered twice daily, and individual refusals were recorded. No hay or concentrate was provided to minimise dietary variation among animals and ensure a standardised nutritional environment, thereby reducing potential confounding effects on physiological, behavioural, and microbiological outcomes.

### Monitoring and clinical evaluation

All calves were examined twice daily by the same veterinary team from arrival (Day -12) until Day +19 post-infection. Clinical assessments included general condition, milk intake, rectal temperature, nasal discharge, coughing, respiratory rate, respiratory effort, and lung auscultation findings.

Clinical scoring was adapted from a previously described scoring system [12]. Rectal temperature was assigned a score of 0 for values below 39.0°C, 1 for values between 39.1 and 40.0°C, 2 for temperatures ranging from 40.1 to 41.0°C, and 3 for temperatures exceeding 41.0°C. Respiratory rate was scored from 0 to 4 according to the following intervals: <35, 35-40, 41-60, 61-80, and >80 breaths per minute, respectively. Nasal discharge and overall clinical condition were each rated as normal (0), mildly affected (1), or markedly affected (2). Dyspnoea severity (absent, mild, moderate, or severe) and reduction in milk intake (0%, <30%, 30-60%, or >60%) were scored on a scale ranging from 0 to 3.

These individual clinical measurements were combined to generate a composite clinical score following the methodology previously described in our team [12]. Calves presenting a score greater than 2 during the morning health assessment were classified as positive for bovine respiratory disease on that particular day. The morning evaluation was selected for disease classification because it preceded any sampling procedures that might influence the clinical variables under observation.

To reduce potential observer-related variation and maintain biosecurity throughout the study, veterinary personnel were assigned exclusively to either challenged or control groups, with no crossover between areas. Two veterinarians conducted clinical assessments independently within each experimental unit, providing multiple observations per animal each day and enabling reliable detection of respiratory disease signs, including cough-associated events.

### Bovine Respiratory Disease severity based on individual clinical score trajectories

To classify animals as mild-, moderate-, or severe-BRDC cases, we used an integrative clustering strategy that combined features derived from clinical scores. BRDC severity category was characterised using an unsupervised *k*-means clustering approach performed in R (v4.6.1, *stats* package). For each animal, variables describing disease burden were computed, including the number of BRDC episodes (defined as uninterrupted periods of positive diagnosis), mean and maximum episode duration, total duration of BRDC positivity, mean and maximum clinical scores, and the total clinical burden expressed as the area under the curve (AUC) of clinical scores. All variables were centred and scaled prior to analysis, and clustering was performed with *k* = 3 clusters, corresponding to the expected biological groups. The *k*-means algorithm was run with multiple initialisations (nstart = 50) and a maximum of 100 iterations to ensure robustness. Cluster interpretation was based on the distribution of variables within each group, allowing assignment of animals according to disease severity.

### Pathogen surveillance and quantification

Prior to inoculation (Day 0), all calves underwent extensive pathogen screening to confirm the absence of major respiratory pathogens. Nasal swabs placed in a medium were analysed by real-time PCR to confirm the absence of *Mannheimia haemolytica*, *Pasteurella multocida*, *M. bovis*, *Histophilus somni*, BCoV, IDV, bovine respiratory syncytial virus (BRSV), and bovine parainfluenza virus type 3 (BPIV-3). The absence of bovine viral diarrhoea virus (BVDV) was verified by ELISA targeting the Erns antigen (IDEXX BVDV Ag/serum, IDEXX Laboratoires, Saint-Denis, France). Serological testing further confirmed that all calves were negative for *M. bovis*-specific antibodies (ELISA, Bio K 302; BioX Diagnostics) and IDV-specific antibodies (hemagglutination inhibition assay). This multi-layered screening ensured that all animals entered the study free of major respiratory pathogens, providing a clean baseline for evaluating the onset of respiratory disease under controlled infection conditions.

Following infection, nasal swabs and faecal swabs were collected on Days +1, +3, +7, +9, +14, and +19 and analysed for *M. bovis*, IDV, and BCoV.

Viral RNA and bacterial DNA were extracted from 170 µl aliquots of nasal swab medium, raw BAL fluid, and faecal swab samples using the NucleoMag Pathogen kit (Macherey-Nagel, Germany) on the KingFisher Flex purification system (Thermo Fisher Scientific, MA, USA) according to the manufacturer’s instructions. The PB1 (polymerase basic 1) gene of IDV, the N (nucleocapsid protein) gene of BCoV, and the oppD (oligopeptide permease D) gene of *M. bovis* were amplified using specific primers and quantified using the iTaq Universal SYBR Green One-Step kit (Bio-Rad Laboratories, USA) on a LightCycler 96 Real-Time PCR System (Roche Diagnostics, Germany). The bovine GAPDH gene was used as an internal control to assess sample quality and the efficiency of nucleic acid extraction.

The following primers were used: IDV PB1 forward (5’-GCTGTTTGCAAGTTGATGGG-3’) and reverse (5’-TGAAAGCAGGTAACTCCAAGG-3’), BCoV N forward (5’-TGGATCAAGATTAGAGTTGGC-3’) and reverse (5’-CCTTGTCCATTCTTCTGACC-3’), and *M. bovis* oppD forward (5’-TCAAGGAACCCCACCAGAT-3’) and reverse (5’-AGGCAAAGTCATTTCTAGGTGCAA-3’).

Copy numbers in samples were determined using standard plasmids containing the PB1 product of D/bovine/France/5920/2014 (IDV), the N product of strain ENVT-66 (BCoV), or the oppD product of strain RM16 (*M. bovis*).

Amplification was performed under the following conditions: reverse transcription at 50 °C for 10 min (IDV and BCoV only), initial denaturation at 95 °C for 1 min, followed by 40 cycles of 95 °C for 15 sec and 60 °C for 40 sec. Each reaction contained 5 µL Master Mix, 0.25 µL of each primer (final concentration 200 nM), 0.125 µL of iScript reverse transcriptase for the IDV and BCoV assays, and nuclease-free water to a final volume of 9 µL, followed by the addition of 1 µL of template.

To estimate the cumulative infectious challenge each calf encountered, we generated an integrated pathogen load index based on quantitative measurements of the three respiratory pathogens under investigation: *Mycoplasma bovis*, BCoV, and IDV in the nasal cavity and in the rectum. Pathogen abundance was assessed using quantitative PCR and expressed as gene copy numbers obtained from both nasal and faecal samples.

Because qPCR-derived gene copy numbers spanned several orders of magnitude and exhibited a strongly right-skewed distribution, all values were transformed using a log₁₀ scale after adding a pseudocount of one copy to account for zero observations. Thus, each measurement was converted according to log₁₀(gene copies + 1).

The transformed values corresponding to the three pathogens were then combined into a single score by summation:

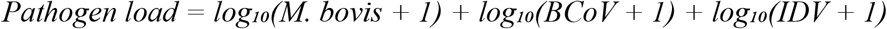

This composite index was designed to represent the overall infectious burden experienced by an individual animal, assuming that the effects of multiple pathogens are additive.

To investigate how pathogen replication dynamics varied over time and among experimental groups, longitudinal analyses were performed together and separately for BCoV, IDV, and *M. bovis*. Because pathogen loads were measured repeatedly in the same animals throughout the study, linear mixed-effects models were used to account for both temporal variation and within-animal correlations. For each pathogen, log₁₀-transformed pathogen load was included as the response variable in a linear mixed-effects model fitted using the lme4 package (v2.0.1) in R. Experimental group, sampling day, and their interaction were specified as fixed effects, while animal identity was included as a random intercept to account for the repeated-measures structure of the dataset. P-values for fixed effects were estimated using Satterthwaite’s approximation implemented in the lmerTest package (v3.2.1). Statistical significance of main effects and Group × Day interactions was assessed using Type III ANOVA. When a significant interaction was detected (*P* < 0.05), post hoc pairwise comparisons between experimental groups were performed at each sampling time point using estimated marginal means (emmeans package v2.0.3). Resulting *P*-values were adjusted for multiple testing using the Benjamini-Hochberg false discovery rate correction.

### Measurement of acute-phase proteins

Serum samples collected from all calves on days −8, −1, +1, +3, +7, +9, +14, and +19 relative to pathogen challenge were analysed to quantify markers of the acute-phase response. This response was evaluated by measuring serum amyloid A (SAA) and haptoglobin (HP). SAA concentrations were quantified using a commercial ELISA assay, whereas HP concentrations were determined using a colourimetric method (Tridelta Development Ltd., Ireland). Absorbance values were recorded with an ELx800 microplate reader (BioTek Instruments, USA) and processed using KC Junior software (BioTek Instruments, USA). All analyses were performed following the protocols provided by the respective manufacturers.

Longitudinal changes in serum SAA and haptoglobin concentrations were analysed using the same linear mixed-effects modelling framework described for pathogen loads. SAA and haptoglobin concentrations were used as the response variable, with experimental group, sampling day, and their interaction included as fixed effects, and animal identity included as a random intercept to account for repeated measurements within calves. Statistical inference, post hoc comparisons, and multiple-testing correction were performed as described above.

### Quantification of Short-Chain Fatty Acids in Faecal Samples

Approximately 1 g of fresh faecal material was mixed with 0.2 mL of 25% (v/v) sulfuric acid to preserve short-chain fatty acids (SCFAs) and prevent microbial degradation. Samples were stored at −20°C until analysis. Prior to measurement, samples were thawed on ice and centrifuged at 2,880 × g for 20 min at 4°C to obtain the liquid fraction. Proteins were then removed by adding 200 µL of 25% (v/v) metaphosphoric acid to 1 mL of supernatant, followed by centrifugation at 20,000 × g for 15 min at 4°C.

For gas chromatographic analysis, 100 µL of the cleared extract was mixed with 75 µL of 0.2% (v/v) 4-methylvaleric acid as an internal standard and diluted with 900 µL of ultrapure water. A 1 µL aliquot was injected into a Hewlett-Packard 7890A gas chromatograph equipped with a flame ionisation detector and a DB-FFAP capillary column. Chromatographic peaks were analysed using Chromeleon software.

We quantified six SCFAs using calibration curves and internal standard normalisation: acetic, propionic, butyric, valeric, isobutyric, and isovaleric acids. Total SCFA concentration was calculated as the sum of all measured acids, while the relative abundance of each SCFA was expressed as a percentage of the total concentration.

To correct for differences in faecal moisture content, SCFA concentrations were normalised to dry matter. Briefly, 5 g of each sample was dried at 60°C for 48 h until a constant weight was achieved. Dry matter percentage was calculated from the ratio of dry to fresh weight, and SCFA concentrations were subsequently expressed per gram of dry faecal matter.

For each SCFA, a separate linear mixed-effects model was fitted using the same analytical framework described for pathogen loads, including experimental group, sampling day, and their interaction as fixed effects, and animal identity as a random effect to account for repeated measurements. Statistical testing and post hoc analyses were performed as described above. For SCFA concentration, the emmtrends() function (Estimation of marginal means of linear trends, emmans R package v. 2.0.3) was used to assess relationships between SCFA concentration and bacterial loads. Moreover, the r2glm R package v. 0.1.3 and the MuMIn R package v. 1.48.9 were used to perform slope confirmatory tests and r² on the linear mixed-effects models. The p-value obtained represents the certainty that the measured slope is non-zero.

### Faecal and nasal microbial DNA extraction

Faecal and nasal swab samples were obtained from all calves at days −8, −1, +1, +3, +7, +9, +14, and +19 relative to experimental infection. Following storage and thawing on ice, total microbial DNA was extracted from faecal samples using the E.Z.N.A.® Stool DNA Kit (Omega Bio-Tek, USA) and from nasal swabs using the ID Gene™ Mag Fast Extraction Kit in combination with the IDEAL 96 automated extraction system (Innovative Diagnostics, France), according to the respective manufacturers’ instructions.

DNA concentration and quality were evaluated using both NanoDrop spectrophotometry (NanoDrop Technologies, USA) and Qubit dsDNA High-Sensitivity fluorometric quantification (Thermo Fisher Scientific, USA). Negative extraction controls were processed alongside all batches to detect potential contamination. To further minimise contamination risk, we used DNA-free reagents and consumables, UV-treated materials, ultrapure water, and physically separated pre- and post-PCR workspaces throughout the workflow.

A standardised microbial mock community (ZymoBIOMICS™ Microbial Community Standard, Zymo Research, USA) was included as a positive extraction control to assess extraction performance and sequencing accuracy. Following quality assessment, all DNA samples were normalised to a concentration of 10 ng/µL before submission for sequencing.

### qPCR-Based Quantification of Nasal and Faecal Bacterial Biomass

Total bacterial biomass in faecal and nasal swabs was quantified by quantitative PCR (qPCR) following the methodology described by Mach’s team [15]. Amplification reactions were performed on a QuantStudio 12K Flex Real-Time PCR System (Thermo Fisher Scientific, Waltham, MA, USA) using universal bacterial 16S rRNA gene primers (forward: 5′-CAGCMGCCGCGGTAANWC-3′; reverse: 5′-CCGTCAATTCMTTTRAGTTT-3′) synthesised by Eurofins Genomics (Ebersberg, Germany).

To enable absolute quantification, purified PCR amplicons generated with the same primer pair were serially diluted ten-fold to construct a seven-point standard curve spanning concentrations from 2.25 × 10⁷ to 2.25 × 10¹³ gene copies per µg of DNA. Bacterial abundance in faecal samples was then estimated by interpolation against this standard curve.

Each qPCR reaction was carried out in a final volume of 20 µL containing 10 µL of Power SYBR Green PCR Master Mix (Thermo Fisher Scientific), 2 µL of template DNA or standard solution (0.5 ng/µL), and primers at a final concentration of 0.6 µM each. Thermal cycling conditions consisted of an initial denaturation step at 95°C for 10 min, followed by 40 amplification cycles comprising 95°C for 15 s and 60°C for 60 s. Amplification specificity was verified by melt-curve analysis performed between 60°C and 95°C at the end of each run. All samples and standards were analysed in triplicate.

Absolute copy numbers were calculated from standard curves generated using purified amplicons. Copy number estimates were derived from DNA mass and amplicon length according to the equation:

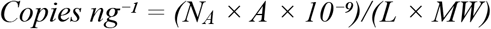

where *N_A_* corresponds to Avogadro’s constant (6.02 × 10²³ molecules mol⁻¹), *A* is the DNA mass (ng), *L* is the amplicon length (bp), and *MW* is the average molecular weight per base pair. To satisfy the assumptions of parametric statistical analyses, bacterial biomass values were log₁₀-transformed prior to analysis. Consequently, results are reported as log₁₀ bacterial gene copies per gram of wet faeces (Supplementary Table S1). Differences in bacterial biomass among experimental groups and sampling time points were subsequently assessed using the same linear mixed-effects modelling framework described above, incorporating animal identity as a random effect to account for repeated measurements. When required, pairwise comparisons were corrected for multiple testing using the Benjamini-Hochberg procedure.

### Faecal and nasal 16S rRNA gene sequencing

The V3-V4 region of the bacterial 16S rRNA gene was amplified from 10 ng of genomic DNA following a previously validated protocol. Library preparation was performed according to Illumina guidelines and automated using a Freedom EVO NGS workstation (TECAN) to reduce contamination and pipetting errors. The V3-V4 region was initially amplified using specific primers and KAPA HiFi HotStart ReadyMix under the following conditions: 95°C for 3 min; 25 cycles of 95°C for 30 s, 55°C for 30 s, and 72°C for 30 s; followed by a final extension at 72°C for 5 min. Amplicons were purified with AMPure XP beads.

Dual indices and Illumina sequencing adapters were subsequently added in a second PCR (8 cycles) using the Nextera XT Index Kit. The resulting libraries were purified, quantified by fluorescence and qPCR, and assessed for quality using an Agilent 2200 TapeStation.

Sequencing was performed by IGA Technology Services (Udine, Italy) on an Illumina MiSeq platform using the MiSeq Reagent Kit v3 (2 × 300 bp paired-end reads).

A commercially available mock microbial community (ATCC® MSA-1003™, containing 20 bacterial strains at defined relative abundances) was included as a sequencing control to monitor sequencing accuracy and error rates. More details on this protocol (such as primer sequences, PCR protocol, quality control, etc.) are available in [15]

### Microbial 16S rRNA sequence processing

Raw paired-end 16S rRNA sequencing reads were quality-checked using FASTQC (v. 0.12.1), and quality reports were consolidated with MultiQC (v. 1.27.1) to assess read quality, sequencing depth, and potential adapter contamination. Primer and adapter sequences were removed using Cutadapt within QIIME 2 (v. 2024.10). High-quality reads were then processed using the DADA2 denoise-paired workflow, which performs quality filtering, error correction, chimaera removal, and inference of amplicon sequence variants (ASVs).

Different DADA2 parameters were applied to faecal and nasal samples to account for sample-specific sequencing characteristics. Taxonomic classification of ASVs was performed using a naïve Bayes classifier trained on the SILVA database (v138-99-nb). Low-abundance ASVs (≤2 reads), as well as mitochondrial and eukaryotic sequences, were removed to improve data quality and retain only prokaryotic microbial signals.

The filtered feature table and taxonomic assignments were imported into R (v. 4.3.1) for downstream analysis. Sequencing accuracy was assessed using mock community controls, while potential contaminants were identified and removed using the decontam package (v. 1.28.0) based on prevalence in negative controls. Subsequent analyses were conducted using phyloseq (v. 1.46.0), vegan (v. 2.6.4), and microbiome (v. 1.23.1) R packages for microbial community integration, diversity assessment, compositional analysis, statistical modelling, and data visualisation.

### Processing and taxonomic classification of microbial sequencing data

Amplicon sequence variant (ASV) tables were analysed using the *phyloseq* package (v1.54.0). To maximise taxonomic resolution, ASVs lacking genus-level annotation were assigned a custom label based on the highest available taxonomic rank, typically combining the term “Unclassified” with the corresponding family designation. Subsequent analyses were conducted using genus-level aggregated data.

To account for differences in sequencing depth among samples, read counts were converted to relative abundances by normalising each sample to its total number of sequences. Summary statistics, including sequencing depth, total read counts, and the number of detected ASVs, were calculated for quality assessment. Animals were classified according to antimicrobial exposure status (ATB vs. no-ATB) for downstream comparisons.

### Within-sample microbial diversity

Microbial richness and diversity were characterised using Observed Richness and Shannon diversity indices computed in *phyloseq*. Temporal changes in alpha diversity were evaluated across sampling points and experimental groups, while accounting for individual pathogen burden. Longitudinal changes in microbial diversity over time and between experimental groups were assessed using the same linear mixed-effects modelling approach described for pathogen loads. Models included group, sampling day, and their interaction as fixed effects, and animal identity as a random effect to account for repeated observations. Statistical testing and multiple-comparison corrections were performed as described above.

### Between-sample diversity and microbial community structure

Microbial community composition was characterised using Bray-Curtis dissimilarity, which accounts for taxon relative abundances, Jaccard distance, which is based solely on taxon presence or absence, and weighted unifrac. Distance matrices were generated using the vegdist() function from the vegan package. We visualised microbial community structure using non-metric multidimensional scaling (NMDS) ordination with the metaMDS() function, allowing comparison of microbiota composition across sampling times, treatment groups, and biological compartments. Group clustering patterns were illustrated by adding confidence ellipses to the ordination plots.

Differences in microbial community composition were tested using permutational multivariate analysis of variance (PERMANOVA) implemented with the adonis2() function (999 permutations). The models included biological compartment (airway versus faecal), sampling time, antibiotic exposure, and total pathogen load as explanatory variables. Statistical significance was determined from permutation-based p-values, and the contribution of each factor to the observed variation was reported as R² values.

To verify the assumptions underlying PERMANOVA, homogeneity of multivariate dispersion among groups was assessed using the betadisper() function, followed by permutation testing with permutest(). This step ensured that any detected differences in community composition were not primarily driven by unequal within-group dispersion.

### Taxonomic Associations with Pathogen Burden and Clinical Severity

To identify bacterial genera associated with treatment, pathogen burden, and clinical outcome, we employed two complementary analytical approaches: Analysis of Compositions of Microbiomes with Bias Correction (ANCOM-BC; v2.14.0) and linear mixed-effects regression models (LMER; lme4 package). The use of both methods enabled robust identification of taxa while accounting for the compositional nature of microbiome data and the longitudinal structure of the experiment.

ANCOM-BC was applied to detect differentially abundant genera while correcting for compositional bias inherent to sequencing datasets. Taxa showing significant differences in abundance across experimental groups, clinical outcome clusters, or pathogen burden categories were identified using bias-corrected log-fold changes and associated statistical tests. In parallel, longitudinal associations between bacterial genera and explanatory variables were assessed using linear mixed-effects models. For each genus, normalised relative abundance was modelled as a function of treatment group, pathogen load, clinical severity cluster, sampling day, and their relevant interactions. Animal identity was included as a random intercept to account for repeated measurements collected from the same calf throughout the study period. To improve robustness and reduce the risk of false-positive discoveries, only taxa detected in a sufficient proportion of samples were retained for analysis. Statistical significance was evaluated independently for each method, and resulting *P*-values were corrected for multiple testing using the Benjamini-Hochberg false discovery rate (FDR) procedure. Genera were considered strongly associated with the outcome of interest when significance was supported by both ANCOM-BC and LMER analyses.

Combining ANCOM-BC and mixed-effects modelling allowed us to identify bacterial taxa consistently associated with disease progression while accounting for compositional effects, temporal dynamics, and repeated observations within individual animals.

## Results

### Antibiotic treatment and experimental inoculation of respiratory pathogens shape BRDC severity, but marked individual variability drives distinct clinical outcomes

To assess bovine respiratory disease complex severity, a veterinarian examined calves twice daily and assigned clinical scores based on multiple respiratory and systemic signs. These scores were integrated into a composite clinical index and used to monitor disease progression throughout the study (Figure 1A). Although we implemented extensive biosecurity measures to prevent pathogen spread between experimental groups, transmission occurred between the coinoculated and non-inoculated pens. Indeed, we experimentally inoculated only half of the calves with *M. bovis*, BCoV, and IDV on Day 0, yet animals housed in the non-inoculated section developed clinical signs compatible with BRDC about three days later. Subsequent pathogen surveillance confirmed cross-transmission within the facility. Specifically, both BCoV and *M. bovis* were detected in calves from all treatment groups, indicating successful circulation despite pen separation and strict biosecurity procedures (Supplementary Table 1). In contrast, IDV remained largely restricted to experimentally inoculated animals, with only sporadic low-level detection in two non-inoculated calves (IDs 27 and 28). Therefore, animals were reclassified according to the route of exposure, distinguishing experimentally infected calves from those that acquired infection naturally through transmission events within the cohort. Combined with antibiotic treatment status, this resulted in four analytical groups: experimentally inoculated and antibiotic-treated (Exp_ATB), experimentally inoculated and untreated (Exp_noATB), naturally infected and antibiotic-treated (Nat_ATB), and naturally infected and untreated (Nat_noATB). We then compared the longitudinal evolution of clinical disease among these four groups.

**Figure 1:**
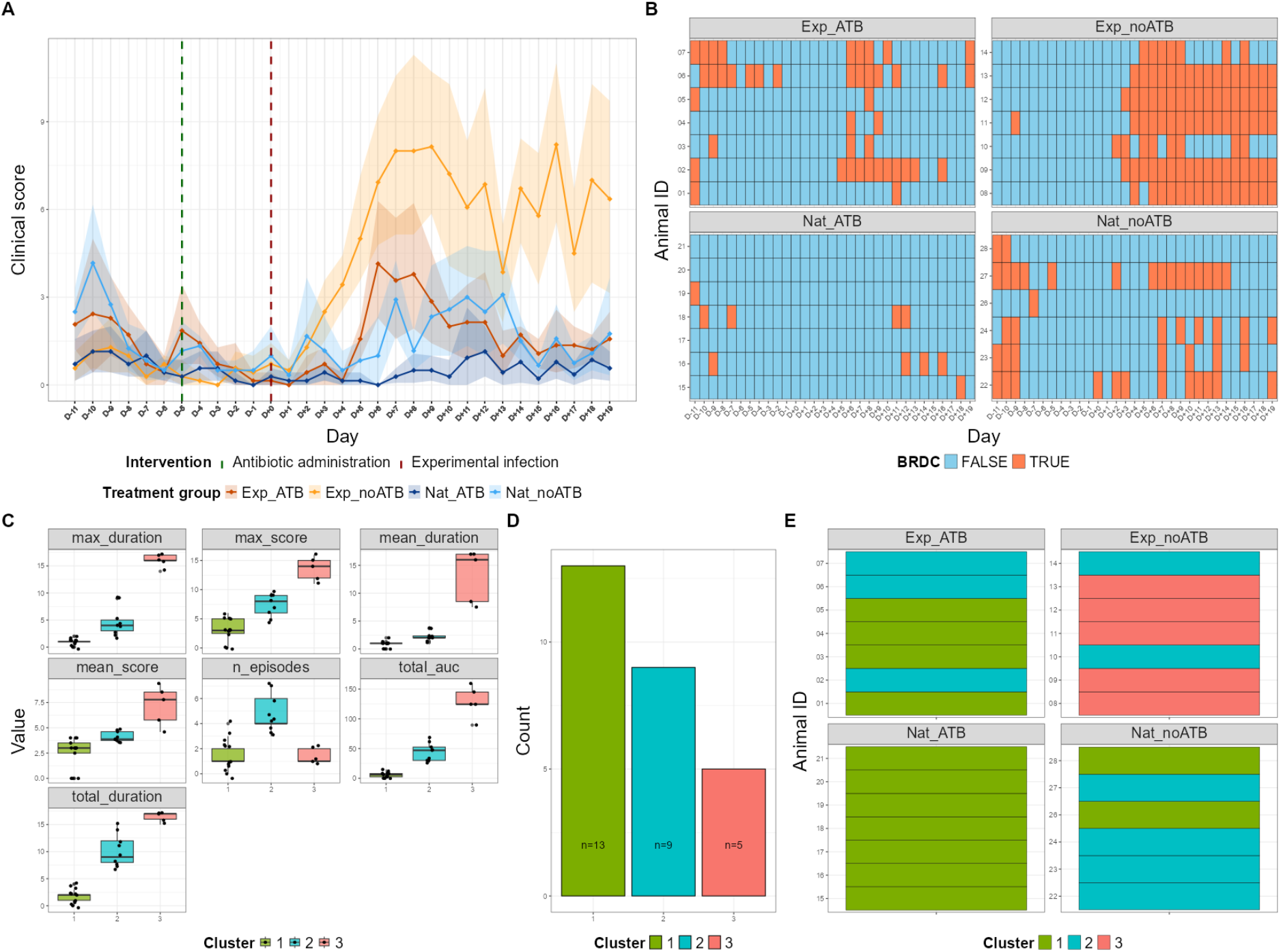
Clinical signs and BRDC severity cluster. **(A)** Mean clinical score per day per experimental group. The green dotted line marks the start of 3-day antibiotic administration. The red dotted line represents the day of experimental exposure to BRDC pathogens. Shaded ribbons represent 95% of the group population. **(B)** Individual Health status per day. If an animal had a clinical score >2 in the morning, it was considered “Positive for BRDC” for the day. **(C)** Distribution of clinical score variables per K-means clustering group. The algorithm used these variables to identify clusters by grouping animals with the same variable profiles. **(D)** Number of individuals in each K-means cluster. **(E)** Individual K-means cluster attribution per experimental condition.

Before pathogen exposure, calves in all groups were clinically healthy, indicating similar baseline health status. Following infection, marked differences in BRDC severity emerged. Calves in the Exp_noATB group consistently exhibited the highest mean clinical scores throughout the study, indicating the most severe clinical manifestations. In contrast, the Nat_ATB group maintained the lowest scores and therefore experienced the mildest disease course. The Exp_ATB and Nat_noATB groups showed similar trajectories, exhibiting intermediate disease severity throughout the study. No consistent divergences were observed between the groups, except for D+6 and D+8 (emmeans post hoc test on LMM, adjusted *P* < 0.05). As the study progressed, clinical scores in both groups gradually declined and converged toward those observed in the Nat_ATB group, reflecting progressive recovery of the animals (Figure 1A).

To complement the group-level analysis, disease occurrence was also examined at the individual level. Calves were classified as BRDC-positive on any day when their morning clinical score exceeded 2 (Figure 1B). This approach enabled us to visualise the timing, duration, and recurrence of disease episodes for each animal throughout the study. Some calves developed severe, persistent clinical signs that remained elevated for extended periods, whereas others had only transient disease episodes followed by rapid recovery. Precisely, nearly all calves experienced at least one BRDC episode, except calves 17, 20, and 21 from the Nat_ATB group, which remained clinically healthy throughout the experiment. Consistent with the mean clinical score trajectories described above, calves in the Nat_ATB group had the lowest disease burden, characterised by fewer and shorter episodes of clinical illness. Conversely, animals that did not receive antibiotic treatment generally experienced prolonged periods of BRDC (Wilcoxon test, W=23, *P* < 0.05).

To objectively characterise this heterogeneity and identify distinct disease-response profiles, we performed an unsupervised *K*-means clustering analysis based on individual clinical score trajectories and BRDC episode characteristics (Figures 1C-E). This approach grouped calves by overall severity, duration, and temporal dynamics of clinical responses, allowing identification of biologically meaningful disease phenotypes beyond the original experimental classifications (Figures 1C–E; Supplementary Table S1). The *K*-means analysis identified three distinct clinical outcome profiles. Cluster 1 corresponded to calves with the mildest disease course, characterised by low clinical scores, short disease episodes, and the lowest cumulative disease burden. Cluster 2 represented an intermediate phenotype, in which calves experienced the greatest number of BRDC episodes, although these episodes were generally transient and less severe. Cluster 3 comprised the most affected animals and was characterised by prolonged periods of illness, elevated clinical scores, and the largest area under the clinical score curve (AUC; Dunn test, *P* < 0.005, Holm adjusted), indicating a substantial cumulative disease burden (Figures 1C-E). Of the 28 calves initially included, one calf was excluded from the clustering analysis because it was identified as an outlier, exhibiting unusually high clinical scores from the beginning of the monitoring period. Among the remaining 27 calves, 13 were assigned to Cluster 1, 9 to Cluster 2, and 5 to Cluster 3.

The distribution of clusters was strongly influenced by both experimental infection status (Fisher’s exact test, *P* < 0.05) and antibiotic treatment (Fisher’s exact test, *P* < 0.005). All calves in the Nat_ATB group belonged to Cluster 1, consistent with the mild disease observed in this group. Similarly, the Exp_ATB group showed no severe cases, with four calves classified as Cluster 1 and three as Cluster 2. In contrast, the Exp_noATB group was dominated by severe outcomes, with five of seven calves assigned to Cluster 3. The Nat_noATB group displayed the greatest heterogeneity, containing animals from all three clusters (two mild, four moderate, and one severe), highlighting the marked variability in host responses following natural pathogen transmission.

### Differential Pathogen Dynamics Underlie Contrasting BRDC Severity Profiles

The marked heterogeneity in clinical severity among calves suggested that factors such as pathogen loads, host immune responses, and microbiota composition and functionality may contribute to individual susceptibility and resilience to BRDC. To test whether disease severity differed by pathogen burden, we first examined the longitudinal dynamics of *M. bovis*, BCoV, and IDV concentrations across experimental groups and disease severity clusters (Figure 2). Prior to pathogen challenge, pathogen loads remained close to zero in all animals (Figure 2A). Following inoculation, a rapid increase in total pathogen burden was observed across all experimental groups, reaching peak values between days 2 and 4 post-challenge (Figure 2A). Animals allocated to the antibiotic-treated groups (Exp_ATB and Nat_ATB) showed lower pathogen loads during the later stages of follow-up than their untreated counterparts (emmeans post hoc test on LMM, estimate =-0.93, *P* < 0.05), although substantial overlap in the 95% confidence intervals indicated considerable inter-individual variability. Analysis of individual pathogens revealed distinct temporal kinetics (Figure 2B). BCoV displayed the earliest and highest replication peak, with gene copy numbers increasing sharply shortly after challenge and declining thereafter (ANOVA type III on LMM, F = 11.26, overall time effect, *P* < 0.05). Unlike BCoV, IDV loads remained relatively low throughout the study and showed only modest fluctuations over time (Dunn’s test, Holm-adjusted *P* < 0.005; overall time effect, *P* < 0.01). *M. bovis* exhibited a delayed pattern compared with BCoV, with bacterial loads increasing after challenge and remaining detectable for a longer period (Dunn’s test, Holm-adjusted *P* < 0.005; overall time effect, *P* < 0.01).

**Figure 2:**
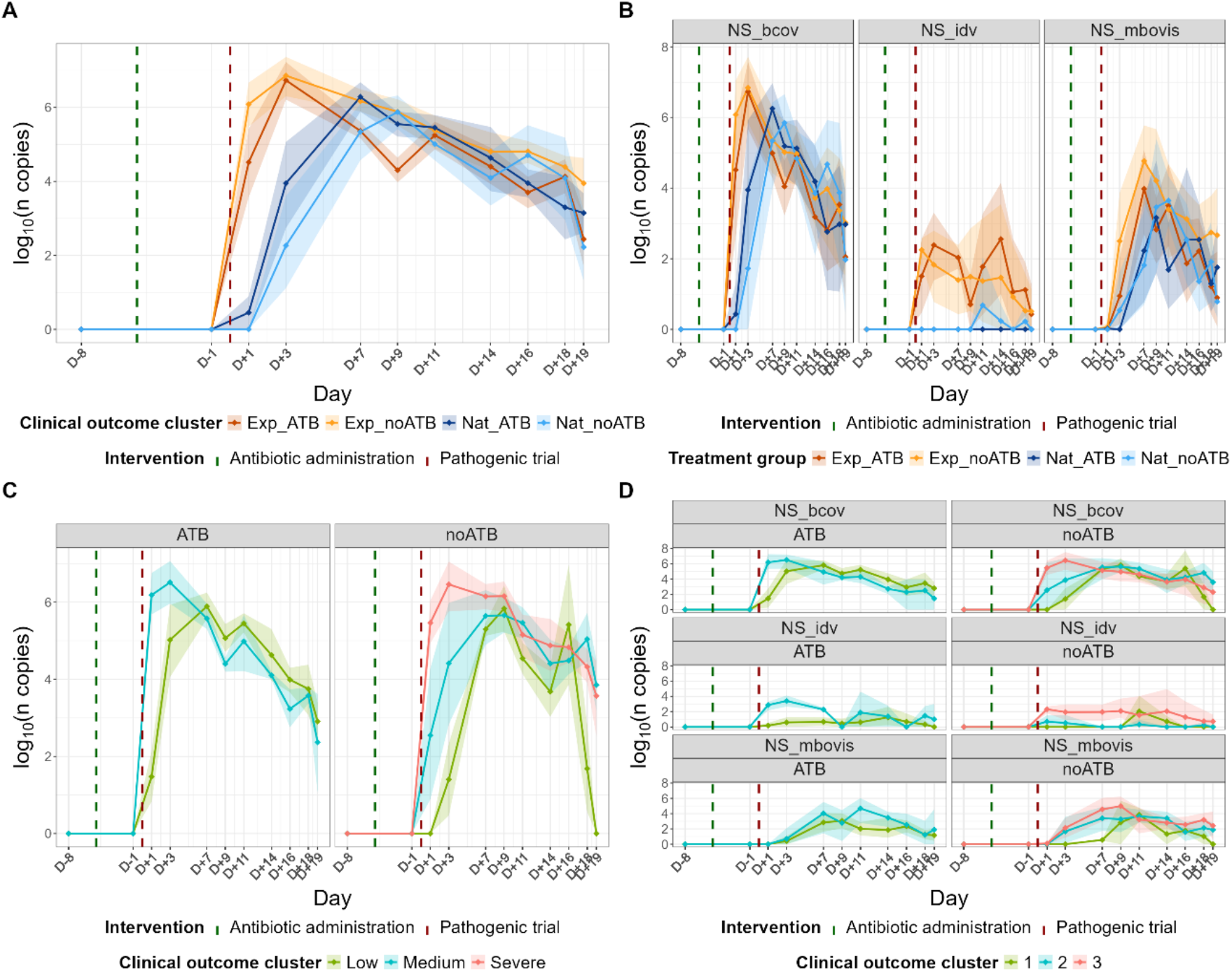
Pathogen loads. **(A)** Total pathogen load per experimental group. Shaded ribbons represent 95% of the group population. **(B)** Mean log10 copies of the targeted qPCR gene for each pathogen of interest per experimental group**. (C)** Total pathogen load per BRDC severity cluster. **(D)** Mean log10 of copies of the targeted qPCR gene for each pathogen of interest per clinical outcome cluster

When animals were stratified according to BRDC severity clusters, total pathogen load differed significantly among severity groups (Figure 2C; type III ANOVA on LMM, cluster effect: *F* = 4.83, *P* < 0.05). In both antibiotic-treated and untreated calves, severity-associated differences were observed at D+1 and D+3 (emmeans post hoc test on LMM, adjusted *P* < 0.05). Thereafter, these differences disappeared in antibiotic-treated animals but persisted in untreated calves, where only the low-severity cluster showed a marked decline in pathogen load at D+18 and D+19 (emmeans post hoc test on LMM, adjusted *P* < 0.05). These findings indicate that severe clinical outcomes were associated with delayed pathogen clearance, particularly in the absence of antibiotic treatment.

Examination of individual pathogen kinetics by clinical severity cluster further supported this association (Figure 2D). Analysis of individual pathogens further showed that this association was primarily driven by IDV, whose abundance differed significantly among BRDC severity clusters (type III ANOVA on LMM, *F* = 4.24, *P* < 0.05). Among untreated calves, severe clinical cases were characterised by persistently elevated IDV loads throughout the acute phase of infection, remaining higher than those observed in less severely affected animals until D+14 (emmeans post hoc test on LMM, adjusted *P* < 0.05; overall cluster effect: type III ANOVA on LMM, *P* < 0.05). In contrast, BCoV loads were higher in severe clinical cases only during the early stages of infection, with significant differences observed up to D+3 (emmeans post hoc test on LMM, adjusted *P* < 0.05). Similarly, *M. bovis* loads differed among severity clusters only transiently, with lower abundances observed in the least severe cluster at D+7 (emmeans post hoc test on LMM, adjusted *P* < 0.05). Collectively, these findings identify sustained IDV replication as the pathogen feature most strongly associated with severe BRDC outcomes in this cohort.

Among the pathogens investigated in rectal swabs, BCoV was the only agent detected in faecal samples. In antibiotic-treated calves, faecal BCoV loads were higher in animals belonging to the more severe clinical clusters at D+7, D+9 and D+14 (emmeans post hoc test on LMM, adjusted *P* < 0.05), suggesting that prolonged enteric shedding of BCoV may also be associated with increased disease severity despite antimicrobial treatment.

To further investigate the mechanisms underlying the observed clinical outcomes, we examined the relationship between pathogen burden and host inflammation using serum amyloid A concentrations as a marker of the acute-phase response. Correlation analyses revealed a weak but significant negative association between pathogen load and SAA concentrations in calves treated with tilmicosin (Supplementary Figure 1; *R*² = 0.08, BH-adjusted *P* < 0.05). Conversely, in untreated calves, particularly those experiencing severe BRDC, pathogen burden tended to be associated with increased SAA concentrations, although this relationship did not reach statistical significance (R² = 0.11, *P* = 0.096, BH-adjusted). This pattern appeared to be driven primarily by IDV, as severely affected untreated calves harboured the highest nasal IDV loads (type III ANOVA on LMM, *P* < 0.05), which coincided with the strongest inflammatory responses. Similar analyses using haptoglobin yielded weaker associations with pathogen burden (Supplementary Figure 2). In both antibiotic-treated and untreated calves, haptoglobin concentrations remained relatively stable across the range of pathogen loads examined, with regression models explaining less than 1% of the observed variation (R² < 0.01). When stratified by disease severity, haptoglobin concentrations tended to increase slightly with pathogen burden in severely affected untreated calves, whereas no clear relationship was observed in low-or moderately affected calves or in animals receiving antimicrobial treatment. However, these trends were weak and highly variable, suggesting that haptoglobin was less responsive to changes in respiratory pathogen burden than SAA in this experimental setting.

Together, these findings suggest that tilmicosin mitigated the clinical expression of BRDC primarily by modulating the host inflammatory response rather than measurably reducing pathogen burden.

### Antibiotic-Induced Dysbiosis Reveals Microbiota Features Associated with Pathogen Burden and BRDC Severity

Given our hypothesis that antibiotic-induced dysbiosis could influence pathogen establishment and disease progression, we next examined whether tilmicosin-induced microbiota perturbations were associated with differences in clinical outcome, pathogen burden, and nasal microbiota structure after challenge. Indeed, we have demonstrated that administration of tilmicosin in these animals successfully generated a dysbiotic state in both the respiratory and intestinal microbiota, characterised by major ecological and functional alterations [55]. Collectively, these findings confirmed that the antibiotic treatment achieved its intended effect of generating a dysbiotic state prior to pathogen exposure.

Having established that antibiotic administration reshaped both airway and gut microbial communities, we next sought to determine whether differences in microbiota structure could help explain the marked heterogeneity in pathogen burden and clinical severity observed following challenge. Importantly, subsequent analyses were performed using quantitative rather than compositional microbiota data. We corrected relative genus abundances derived from 16S rRNA gene sequencing using total bacterial biomass (16S rRNA gene copy numbers) measured in each sample, allowing estimation of the absolute abundance of individual taxa.

At the level of alpha diversity, however, neither the nasal nor the faecal microbiota showed a clear or consistent association with BRDC severity (Figures 3–4). In the nasal microbiota, only limited and transient differences were observed among severity clusters, involving Pielou’s evenness in untreated calves and richness at isolated time points in antibiotic-treated animals (emmeans post hoc test on LMM, adjusted *P* < 0.05; Figure 3A). However, no overall effect of disease severity on nasal alpha diversity was detected (type III ANOVA on LMM, *P* > 0.05). Similarly, BRDC severity was not consistently associated with faecal microbial richness, evenness, or Shannon diversity in either treatment group (Figure 4A; type III ANOVA on LMM, *P* > 0.05), although occasional time-specific differences were observed (emmeans post hoc test on LMM, adjusted *P* < 0.05).

**Figure 3:**
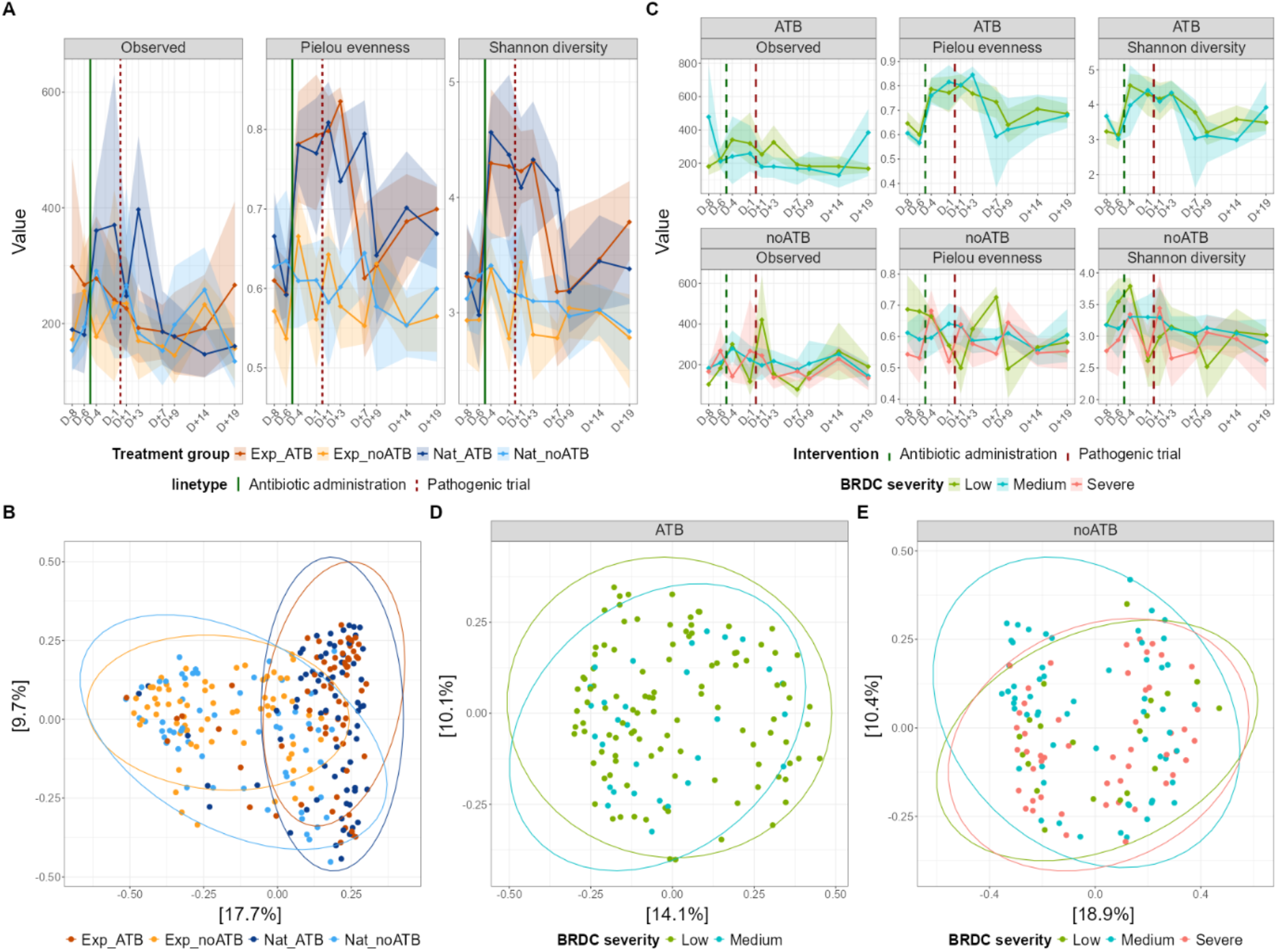
Nasal microbiota alpha and beta diversity and composition stability. **(A)** Mean alpha diversity values per experimental group. Shaded ribbons represent 95% of the group population. **(B)** PCoA of Bray-Curtis dissimilarities per experimental group. **(C)** Mean alpha diversity values per BRDC severity cluster. Shaded ribbons represent 95% of the cluster group population. **(D)** PCoA of Bray-Curtis dissimilarities per BRDC severity cluster, for groups treated with an antibiotic. **(E)** PCoA of Bray-Curtis dissimilarities per BRDC severity clusters, for groups not treated with antibiotic.

**Figure 4:**
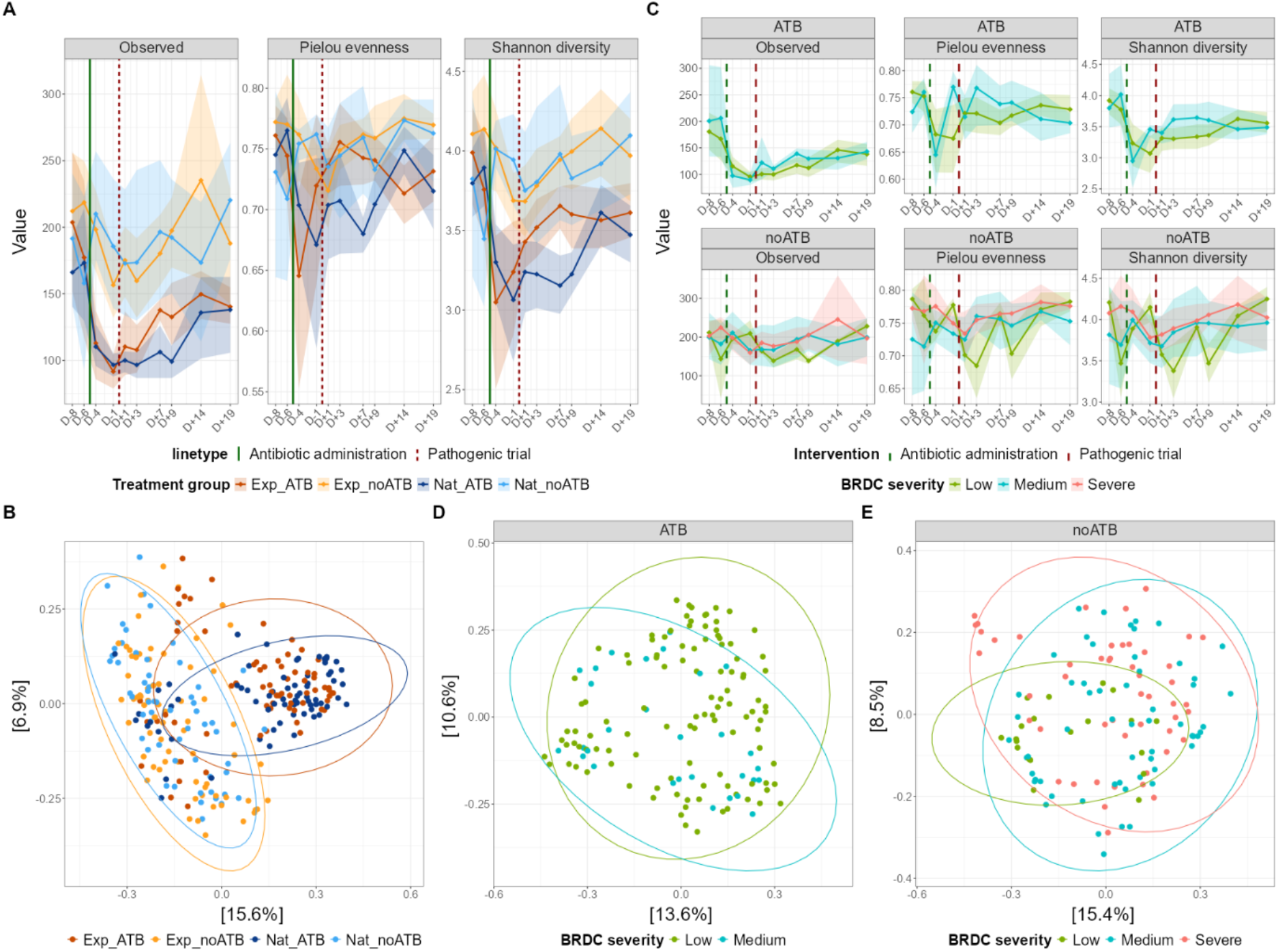
Faecal microbiota alpha and beta diversity and composition stability. **(A)** Mean alpha diversity values per experimental groups. Shaded ribbons represent 95% of the group population. **(B)** PCoA of Bray-Curtis dissimilarities per experimental group. **(C)** Mean alpha diversity values per BRDC severity clusters. Shaded ribbons represent 95% of the cluster group population. **(D)** PCoA of Bray-Curtis dissimilarities per BRDC severity clusters, for groups treated with antibiotic. **(E)** PCoA of Bray-Curtis dissimilarities per BRDC severity clusters, for groups not treated with antibiotic.

Despite the profound respiratory dysbiosis induced by tilmicosin in this cohort [55], analysis of the nasal microbiota revealed no significant relationship between microbial community structure and BRDC severity, regardless of whether compositional differences were assessed using Bray-Curtis or Weighted UniFrac distances (Figures 3D and 3E; PERMANOVA, in all cases, *R*² < 0.04, *P* > 0.01). Thus, although antibiotic administration markedly reshaped the nasal microbiota, the resulting dysbiotic communities did not segregate by disease severity and failed to explain heterogeneity in clinical outcomes. Animals in the mild, moderate, and severe clinical clusters showed largely overlapping nasal microbial communities throughout the study (Figure 3D-E), indicating that variation in nasal community structure after antibiotic exposure did not drive BRDC severity. Interestingly, this pattern contrasted with observations made before induction of dysbiosis (Supplementary Figure 3). Analysis of the pre-treatment nasal microbiota (D-8 and D-6) revealed significant differences among BRDC severity groups (PERMANOVA, *R*² = 0.08, *P* < 0.05). Calves that subsequently developed severe disease exhibited a more homogeneous nasal microbiota than calves with mild or moderate disease (Bray-Curtis distance; PERMANOVA, *R*² = 0.07865, *P* < 0.05; PERMDISP, *P* < 0.05; Tukey’s HSD, *P* < 0.015), suggesting that susceptibility to severe BRDC may be linked to pre-existing nasal microbiota features present before infection. However, once antibiotic-induced dysbiosis was established, these baseline differences disappeared (Bray-Curtis distance; PERMANOVA, *P* > 0.05).

In contrast, the intestinal microbiota displayed the opposite pattern. While pre-existing nasal community structure was associated with future disease severity, it lost its discriminatory power after antibiotic exposure; antibiotic-induced gut dysbiosis, by contrast, was accompanied by clear severity-associated shifts in microbial community structure. Among antibiotic-treated calves, faecal microbiota composition was significantly associated with BRDC severity when assessed using both Bray-Curtis and Weighted UniFrac distances (Bray-Curtis PERMANOVA, *R*² = 0.06, *P* < 0.05; Weighted UniFrac PERMANOVA, *R*² = 0.02, *P* < 0.05; Supplementary Figure 4). More specifically, calves exhibiting moderate clinical severity showed significantly lower distances to the group centroid than calves with mild or severe disease, indicating greater microbiota homogeneity across individuals (Jaccard, Bray-Curtis, and Weighted UniFrac distances; emmeans post hoc test on LMM, adjusted *P* < 0.05). Despite these compositional differences, microbiota drift from baseline was generally comparable among severity groups, with the exception of D+3, when a transient difference was detected using the Bray-Curtis metric (Supplementary Figure 4; emmeans post hoc test on LMM, adjusted *P* < 0.05). Taken together, these results suggest that the early-days nasal microbiota may be relevant for investigating BRDC severity. The faecal microbiota, by contrast, also appears linked to BRDC severity, but only in a dysbiosis context.

### Respiratory and intestinal Taxonomic and functional signatures associate with BRDC severity

Having confirmed that our experimental intervention successfully generated the dysbiotic state predicted by our hypothesis, we next investigated whether specific bacterial taxa and their metabolic outputs could explain differences in pathogen burden and BRDC severity.

Using a combination of linear mixed-effects models and ANCOM-BC, we identified bacterial genera associated with BRDC severity in both faecal and nasal microbiota and subsequently examined their relationships with systemic inflammation and SCFAs.

The strongest and most consistent associations were observed in the faecal microbiota of calves treated with tilmicosin (Figure 5A). Several genera differed between mild and moderate BRDC phenotypes in treated animals, including *Bifidobacterium*, *Lactobacillus*, *Limosilactobacillus*, *Atopobium*, *Collinsella*, and *Megasphaera* (Supplementary Tables S2). However, after correction for multiple testing, only *Chlamydia* remained significant. Importantly, ANCOM-BC also independently identified *Chlamydia*, making it the most robust taxonomic marker associated with BRDC severity in antibiotic-treated calves. In untreated calves, several faecal genera were similarly associated with BRDC severity (Figure 5B), although none remained significant after multiple-testing correction or were independently validated by ANCOM-BC (Supplementary Table S3). Interestingly, the relationship between *Chlamydia* abundance and disease severity appeared opposite to that observed in antibiotic-treated animals, with higher abundances associated with reduced disease severity.

Compared with the gut microbiota, relatively few severity-associated taxa were detected within the nasal microbiota (Supplementary Table S4). Among untreated calves, genera including *Bifidobacterium*, *Ligilactobacillus*, and *Olsenella* showed associations with BRDC severity, but none survived false discovery rate correction.

To assess the biological relevance of these taxa, we then examined their relationships with systemic inflammatory biomarkers. Several faecal genera were associated with SAA concentrations in untreated calves (Supplementary Table S5 and Supplementary Figure 5). Notably, the *Ruminococcus gauvreauii group* displayed contrasting relationships across BRDC severity clusters. In calves with mild disease, this genus was positively associated with SAA concentrations (slope = 0.685, adjusted *P* = 0.005), whereas in the moderate-severity cluster it showed a strong negative association with SAA (slope = -12.6, adjusted *P* = 2.35 × 10⁻⁴²). In untreated animals, the strongest associations were in the *Eubacterium fissicatena group* in the low-severity cluster and *in Chlamydia* in the moderate-severity cluster, suggesting that different microbial taxa may be linked to the inflammatory response at different disease stages or severities. While some genera, such as *Alloprevotella*, were positively associated with SAA concentrations, others, including *Escherichia-Shigella*, showed the opposite pattern (Supplementary Table 5).

Within the nasal microbiota of untreated calves, the strongest association was observed for *Bifidobacterium* in the low-severity cluster, where abundance increased with increasing SAA concentrations (*R*² = 0.41, slope = 1.33, adjusted *P* = 1.3 × 10⁻⁵). A similar pattern was observed for *Olsenella,* which was positively associated with SAA in both low-(*R*² = 0.45, slope = 0.45, adjusted *P* = 0.025) and severe-severity calves (*R*² = 0.13, slope = 0.19, adjusted *P* = 0.008), but negatively associated with SAA in the moderate-severity cluster (*R*² = 0.12, slope = -0.34, adjusted *P* = 0.025). Likewise, *Ligilactobacillus* displayed contrasting relationships across severity groups, being negatively associated with SAA in moderately affected calves (*R*² = 0.15, slope = -0.21, adjusted *P* = 0.008) and positively associated with SAA in severe cases (*R*² = 0.10, slope = 0.19, adjusted *P* = 0.067). Together, these findings indicate that associations between the nasal microbiota and systemic inflammation were heavily dependent on BRDC severity, with several taxa showing opposite relationships with SAA across clinical outcome groups (Supplementary Table 6).

Because several of the bacterial genera associated with BRDC severity are known fermentative taxa, we next investigated whether these microbial signatures were linked to alterations in faecal SCFA concentrations. Before assessing taxa-SCFA associations, we first evaluated the effects of antibiotic treatment and pathogen exposure on the faecal metabolite profile. Tilmicosin treatment significantly altered SCFA concentrations, leading to lower levels of propionate (estimate = -1.84, *P* = 0.0017), valerate (estimate = -0.31, *P* = 0.0037), isobutyrate (estimate = -0.29, *P* = 0.0144), and isovalerate (estimate = -0.33, *P* = 0.0126) relative to untreated calves (Supplementary Table 7). In contrast, pathogen exposure had no detectable effect on any of the measured SCFAs, including acetate, propionate, butyrate, isobutyrate, valerate, and isovalerate (all *P* > 0.05; Supplementary Table 7). Consistent with these observations, no significant associations were detected between nasal bacterial taxa and faecal SCFA concentrations, suggesting that variation in SCFA profiles was driven primarily by changes in the intestinal microbiota. We therefore focused subsequent analyses on relationships between severity-associated faecal genera and SCFA concentrations. In untreated calves, however, only a limited number of significant associations were identified (Table 1). In antibiotic-treated calves, several genera previously associated with milder disease outcomes were positively associated with SCFAs known to arise from bacterial fermentation. Notably, *Bifidobacterium* was positively associated with propionate and valerate concentrations, while *Megasphaera* was associated with isobutyrate and valerate concentrations, particularly valerate production (*R*² = 0.27, adjusted *P* < 0.001). We also detected severity-dependent associations for *Atopobium* and *Lactobacillus*, suggesting that the metabolic consequences of bacterial abundance differ by disease outcome (Table 1).

**Table 1.**
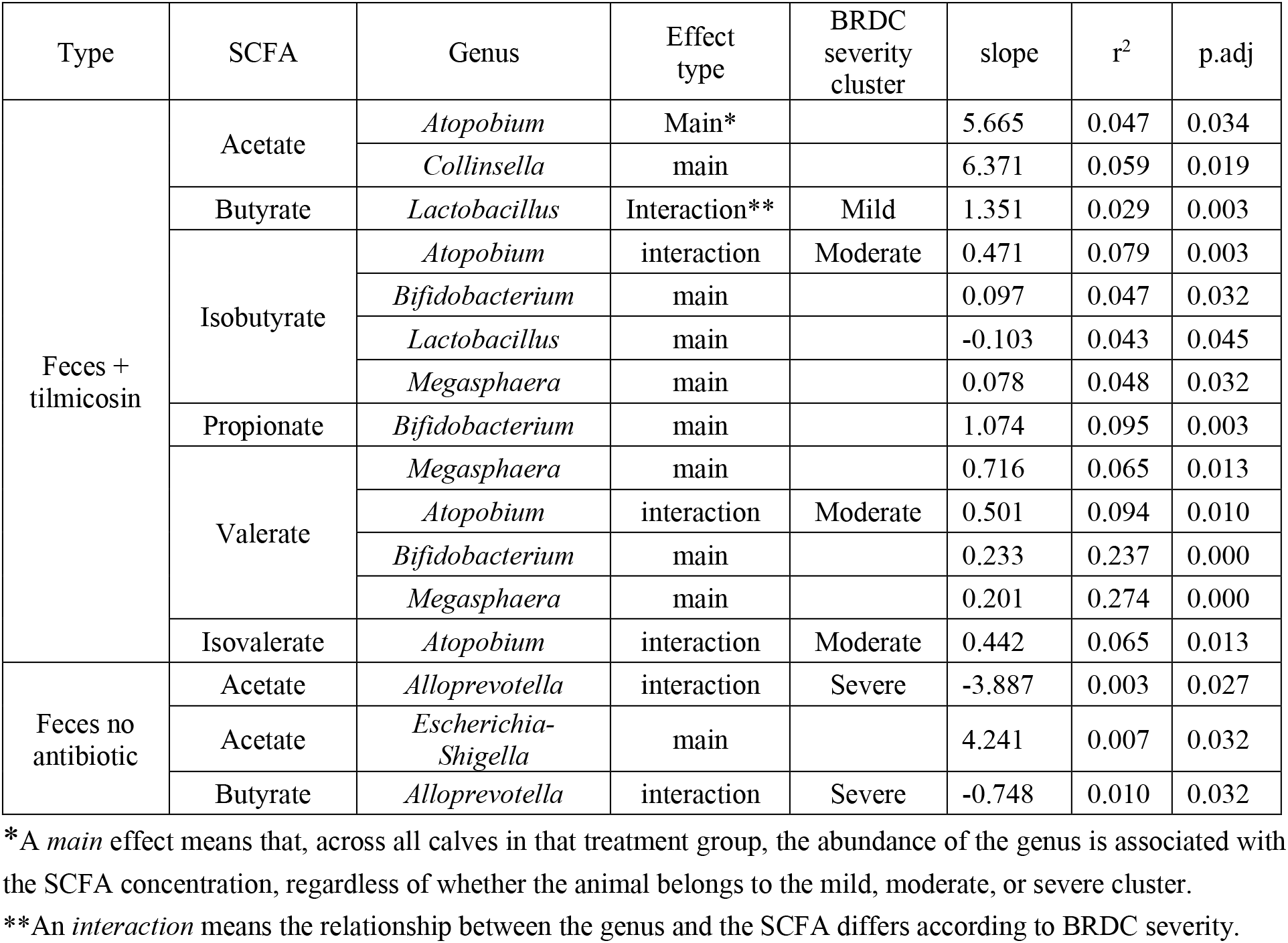
Results of SCFA faecal concentration as a function of bacterial genus abundance and BRDC severity.

In contrast, far fewer associations were detected in untreated calves. The most notable pattern involved *Alloprevotella*, a genus associated with BRDC severity, which was negatively associated with acetate and butyrate concentrations in severely affected animals *(in all cases, R*² < 0.01, adjusted *P* < 0.05). Additionally, *Escherichia-Shigella* abundance was positively associated with acetate concentrations (*R*² = 0.007, adjusted *P* < 0.03). Overall, the limited number of taxa-SCFA associations observed in untreated animals suggests that disease severity was linked primarily to specific microbial taxa rather than broad alterations in fermentative activity, whereas antibiotic treatment appeared to strengthen the relationship between microbial composition and metabolic output. Together, these results indicate that BRDC severity is associated not only with shifts in specific gut bacterial taxa but also with corresponding changes in microbial metabolic activity.

## Discussion

The bovine respiratory disease complex is a multifactorial syndrome resulting from intricate interactions among viral and bacterial pathogens, the host immune system, environmental stressors, and the resident microbiota [1, 14, 17, 59].

This study shows that calves exposed to a similar respiratory pathogen challenge can have markedly different clinical outcomes, and that pathogen burden alone explains only part of this variability. Using a model in which antibiotic-induced dysbiosis was experimentally generated prior to infection, we found that (i) severe BRDC was associated with persistent IDV infection and heightened systemic inflammation, (ii) pre-existing nasal microbiota characteristics were associated with subsequent disease susceptibility, (iii) antibiotic-induced dysbiosis abolished these baseline nasal microbiota signatures, and (iv) alterations in the intestinal microbiota and its metabolic activity were more consistently associated with disease severity than changes in the respiratory microbiota. Together, these findings support the hypothesis that microbiota disruption influences BRDC outcomes and highlight the gut microbiota as a potential contributor to respiratory disease resilience.

Our results should be interpreted in the context of our previous work in the same cohort, which demonstrated that tilmicosin administration successfully induced profound respiratory and intestinal dysbiosis characterised by increased community drift, decreased stability, and marked reductions in short-chain fatty acid concentrations (Mendez et al., 2026)[55]. Building on these observations, the present study shows that the consequences of dysbiosis extend beyond microbial community structure and are associated with differences in clinical outcomes following respiratory infection. Importantly, these effects occurred despite similar pathogen inoculation, indicating that disease severity was not simply driven by pathogen presence.

Consistent with this interpretation, tilmicosin administration did not significantly affect nasal loads of any of the challenge pathogens, including BCoV, IDV, and *M. bovis*. While the absence of an effect on BCoV and IDV was expected because antibiotics lack antiviral activity, the lack of effect on *M. bovis* reflects the high resistance of the challenge strain, as previously reported [55]. Consequently, the reduction in disease severity observed in treated calves cannot be attributed to direct suppression of the inoculated pathogens. Instead, these findings suggest that antibiotic administration modified host-pathogen interactions through mechanisms independent of pathogen clearance, likely due to changes in microbial ecology and function, immune regulation, or both. Thus, one possible explanation may involve the immunomodulatory properties of macrolides. Beyond their antimicrobial activity, macrolides can regulate inflammatory signaling pathways and modulate leukocyte function [60]. The lower circulating SAA concentrations observed in tilmicosin-treated calves, together with the negative relationship between pathogen burden and SAA in treated animals, suggest a more controlled inflammatory response following pathogen inoculation [56]. Although our study was not designed to disentangle microbiota-mediated and immunomodulatory effects of tilmicosin, these findings indicate that both mechanisms may have contributed to the reduced clinical severity observed in treated calves.

Severe BRDC cases occurred exclusively in untreated calves despite broadly similar IDV shedding dynamics between treatment groups. Nevertheless, calves that developed the most severe clinical signs tended to maintain elevated IDV loads over time, suggesting persistent rather than transient infection. Such sustained infections could promote disease progression through alterations of respiratory epithelial function, disruption of mucosal homeostasis, and modulation of immune responses [12, 61, 62]. These observations align with previous studies showing that the most severe BRDC outcomes frequently arise from synergistic interactions between viral and bacterial pathogens [14, 63].

One of the most intriguing findings was the different behaviour of the respiratory and intestinal microbiota. Although tilmicosin successfully induced dysbiosis in both ecosystems, only the intestinal microbiota exhibited clear associations with BRDC severity following antibiotic exposure. In contrast, neither alpha diversity nor overall nasal community composition during infection discriminated calves with mild, moderate, or severe disease. Thus, despite substantial perturbation of the respiratory microbiota, the resulting dysbiotic communities did not segregate according to disease outcome. The biological significance of greater community homogeneity remains uncertain. Unlike previous studies that linked reduced diversity to BRDC susceptibility [18] and risk to disease development later in life [43], our findings point to decreased inter-individual variability rather than reduced within-animal diversity. One possibility is that more homogeneous microbial communities reflect reduced ecological flexibility, limiting the capacity of the respiratory microbiota to respond to infectious challenges and maintain homeostasis. Conversely, greater inter-individual variability may indicate the presence of a broader range of alternative community configurations capable of maintaining ecosystem function following perturbation, a concept consistent with ecological theories of microbiome resilience and alternative stable states [64]. In a opposite way, faecal microbial community structure was significantly associated with BRDC severity in antibiotic-treated animals, supporting our initial hypothesis that disruption of gut microbial ecology may alter susceptibility to respiratory disease [17, 46]. Notably, these associations emerged only after dysbiosis induction and were not observed in the respiratory microbiota, suggesting that the consequences of microbiota disruption differ substantially between mucosal compartments. The taxonomic analyses further reinforced this interpretation. While relatively few nasal taxa were associated with disease outcome, multiple faecal genera were linked to BRDC severity and, importantly, many of these taxa were associated with SCFA concentrations. This pattern suggests that microbial function may be more relevant than taxonomy *per se* [65]. Rather than identifying a single pathogenic or protective bacterial group, our results point toward altered microbial metabolic capacity as a key feature associated with disease outcome.

Although only *Chlamydia* remained significant after correction for multiple testing, the broader ecological pattern was remarkably coherent. Genera associated with milder disease were generally linked to higher SCFA concentrations and lower inflammatory markers, whereas genera associated with severe disease tended to show the opposite relationships. These observations suggest that microbial fermentation capacity and its metabolic products may contribute to regulating host responses during respiratory infection [66]. Particularly noteworthy were the dynamics observed in antibiotic-treated calves. *Bifidobacterium* and *Megasphaera* increased after the peak of clinical signs and were positively associated with valerate concentrations. This temporal pattern suggests that these taxa may be involved in recovery rather than protection against infection itself. Such findings support a model in which the gut microbiota contributes to restoration of host homeostasis following respiratory challenge, potentially through production of metabolites capable of modulating immune and inflammatory processes.

The observations in untreated calves were consistent with this interpretation. *Alloprevotella*, the only genus consistently associated with more severe disease, showed negative correlations with acetate and butyrate concentrations, whereas *Escherichia-Shigella*, which was associated with milder disease, positively correlated with acetate. Although these associations do not establish causality, they suggest that calves developing severe BRDC may have a reduced capacity to maintain microbial fermentation and SCFA production during infection.

Collectively, our findings support a model in which disease severity is shaped by interactions between pathogen persistence, host inflammatory responses, and microbiota-derived metabolic functions. Calves exposed to similar pathogen burdens may therefore experience markedly different outcomes depending on their ability to maintain microbial ecosystem stability and preserve beneficial metabolic activity during infection. The association between severe disease, persistent IDV infection, elevated inflammatory markers, intestinal dysbiosis, and reduced SCFA concentrations provides a coherent framework linking these observations.

## Conclusions

Taken together, our findings demonstrate that antibiotic-induced dysbiosis influences the relationship between pathogen exposure and BRDC outcome. Although tilmicosin administration did not alter pathogen loads, it modified disease severity, indicating that pathogen burden alone is insufficient to explain the marked heterogeneity in clinical responses observed among calves. Instead, our results suggest that disease outcome is shaped by complex interactions among pathogen persistence, host inflammatory responses, and microbiota-associated functions.

Importantly, we identified pre-existing nasal microbiota characteristics associated with subsequent disease susceptibility, whereas antibiotic-induced respiratory dysbiosis largely abolished these baseline microbial signatures. In contrast, dysbiotic intestinal communities remained strongly associated with BRDC severity and were linked to alterations in SCFA production and systemic inflammation. These findings indicate that the consequences of microbiota disruption differ across mucosal ecosystems and point to a more prominent role of the intestinal microbiota in determining disease resilience.

Overall, this study supports the existence of a functional gut-lung axis in calves and highlights microbial metabolic activity as a potential determinant of respiratory health. Rather than being solely a consequence of infection, microbiota composition and function may actively contribute to host susceptibility and recovery following respiratory challenge. Future studies integrating microbiome, metabolomic, and immunological approaches will be needed to establish causality and identify microbial targets for interventions aimed at improving BRDC resilience and reducing dependence on antimicrobials in cattle production systems.

## Limitations of the study

Several limitations should be considered when interpreting these findings. First, the study design introduced an inherent confounding effect between experimental pathogen inoculation, natural pathogen transmission, and antimicrobial treatment. While this feature better reflects field conditions, it also complicates the interpretation of pathogen-specific effects and limits the ability to attribute observed responses exclusively to experimental inoculation. Another important limitation is that pathogen monitoring was restricted to *Mycoplasma bovis*, BCoV, and IDV. Although these pathogens are recognised contributors to BRDC and were the primary targets of the challenge model, the bovine respiratory disease complex is inherently multifactorial. Some variation in clinical severity, inflammatory responses, and microbiota composition may have been influenced by unmeasured pathogens or pathogen-pathobiont interactions.

Third, the sample size, while comparable to other longitudinal large-animal challenge studies, may have limited the detection of weaker associations within the high-dimensional microbiome, pathogen, and host-response datasets. Consequently, several taxonomic associations should be regarded as exploratory and require validation in independent cohorts. Fourth, our microbiota analyses relied on 16S rRNA gene sequencing, which provides taxonomic resolution but limited insight into microbial functionality. Although SCFA measurements provided complementary functional information, confirming the biological mechanisms underlying the observed associations would require direct assessment of microbial metabolic pathways through metagenomics, metatranscriptomics, or metabolomics.

Fifth, the relatively low number of nasal taxa associated with BRDC severity may reflect the respiratory microbiota’s inherently dynamic nature, characterised by high temporal variability and transient colonisation. As a result, potentially relevant microbial signals may have been more difficult to detect than those occurring within the more stable intestinal ecosystem. In addition, the stronger associations observed in the faecal microbiota should not be interpreted as evidence that the respiratory microbiota is unimportant for BRDC pathogenesis, but rather that its contribution may be mediated through ecological processes not fully captured by 16S-based approaches.

Finally, the observational nature of the microbiota, SCFA, and inflammatory analyses precludes causal inference. Whether the identified taxa and metabolites actively influence respiratory disease susceptibility, or instead represent biomarkers of broader host-microbiota interactions, remains unknown. Controlled intervention studies targeting specific microbial functions, metabolites, or microbial communities will be necessary to establish causality and determine their potential for BRDC prevention. Despite these limitations, the longitudinal design, controlled pathogen challenge, simultaneous investigation of respiratory and intestinal microbial ecosystems, and integration of pathogen, microbiota, inflammatory, and metabolic data provide a comprehensive framework for exploring host-microbiota-pathogen interactions during BRDC.

## Data Availability declaration

16S rRNA gene sequences from the nasal cavity and faeces are available in the DDBJ/EMBL/GenBank under BioProject ID PRJNA1517313 at the NCBI Sequence Read Archive (SRA) (www.ncbi.nlm.nih.gov/sra/PRJNA1517313). The temporary accession submission ID is SUB16425776. The associated metadata for this project are available in the INRAE institutional data repository, powered by Dataverse, and are publicly available as of the date of publication (DOI: https://doi.org/10.57745/QJFTEX). The lead contact can provide any additional information required to reanalyse the data reported in this paper upon request.

## Acknowledgments

We sincerely thank the postdoctoral researchers, technicians, and veterinary students who ensured the animals’ daily care and welfare throughout the experiment. In particular, we are grateful to Corto Martin-Rebouissou, Romain Bonsergent, Valentin Puech, Christelle Camus and Chloé Saada for their dedication and support during the study. We also thank Herbipole infrastructure for technical support.

We thank the MIGALE bioinformatics facility for providing support, computing, and storage resources (https://doi.org/10.15454/1.5572390655343293E12; Jouy-en-Josas, France).

## Author Contribution

MMI participated in the experimental study, drafted the manuscript, performed bioinformatic processing of the 16S rRNA sequencing data and biostatistical analyses, integrated the data, and generated most of the figures. TP participated in the experimental study and performed the pathogen-specific PCR assays. GD participated in the experimental study. MC conducted the clinical examinations of the animals during the experimental study. FM participated in the experimental study. DBC contributed to the experimental study. FZS prepared the BCoV and IDV inocula and assisted with the experimental work. CC, XN, and EB provided critical feedback on the manuscript. EB prepared the *M. bovis* inoculum, calculated its antibiotic resistance, and isolated *M. bovis*. XN performed the whole-genome sequencing analysis of the *M. bovis* isolate. GM and MD supported NM in project conception, funding acquisition, and overall project direction. NM helped to draft the manuscript and supervised the bioinformatics and biostatistical analysis.

## Supplementary Tables

**Supplementary Table 1: Metadata and phenotypic characteristics of calves included in the study.** This table contains the metadata, clinical measurements, inflammatory biomarkers, microbiological variables, and metabolite concentrations used to characterise calves throughout the study. Variables include sampling day, animal identification, experimental group, demographic characteristics (birth date, birth weight, sex, and breed), clinical disease score, faecal dry matter content, total nasal bacterial load (16S rRNA gene copies), pathogen loads, faecal short-chain fatty acid (SCFA) concentrations, and serum inflammatory biomarkers including serum amyloid A (SAA) and haptoglobin (HP). Disease severity clusters were derived from longitudinal clinical score trajectories. The variable cluster indicates the numerical cluster assignment generated by the clustering algorithm, whereas cluster_labels indicates the corresponding clinical severity category (Low, Moderate, or Severe).

**Supplementary Table 2: Differentially abundant faecal bacterial taxa associated with BRDC severity in antibiotic-treated calves using ANCOM-BC2.** For each genus, the table reports the omnibus LRT test statistic (compared to a Chi-square distribution), the estimated model coefficient for the moderate BRDC cluster relative to the reference cluster, the omnibus LRT *P*-value, and the Benjamini-Hochberg (BH)-adjusted *P*-value. Positive coefficient values indicate higher abundance in the moderate severity cluster compared with the reference cluster, whereas negative values indicate lower abundance. The omnibus ANOVA tests whether abundance differs across BRDC severity groups overall. Adjusted *P*-values account for multiple testing.

**Supplementary Table 3: Differentially abundant faecal bacterial taxa associated with BRDC severity in non-antibiotic-treated calves using ANCOM-BC2.** For each genus, the table reports the omnibus LRT test statistic (compared to a Chi-square distribution), the estimated model coefficient for the moderate and severe BRDC cluster relative to the reference cluster, the omnibus LRT *P*-value, and the Benjamini-Hochberg (BH)-adjusted *P*-value. Positive coefficient values indicate higher abundance in the moderate severity cluster compared with the reference cluster, whereas negative values indicate lower abundance. The omnibus ANOVA tests whether abundance differs across BRDC severity groups overall. Adjusted *P*-values account for multiple testing.

**Supplementary Table 4. Differentially abundant nasal bacterial genera associated with BRDC severity in non-antibiotic-treated calves.** Differential abundance analysis of the nasal microbiota was performed using ANCOM-BC2. For each bacterial genus the table reports the omnibus LRT test statistic (compared to a Chi-square distribution), the estimated regression coefficient for the moderate and severe BRDC cluster relative to the reference cluster, the omnibus LRT *P*-value, and the Benjamini-Hochberg (BH)-adjusted *P*-value. The omnibus ANOVA tests whether genus abundance differs across BRDC severity groups (Low, Moderate, and Severe), whereas the regression coefficient indicates the direction and magnitude of the abundance change for the corresponding severity cluster relative to the reference level. Positive coefficients indicate higher relative abundance and negative coefficients indicate lower relative abundance compared with the reference cluster. Adjusted *P*-values account for multiple testing using the BH procedure.

**Supplementary Table 5. Associations between faecal bacterial genera and serum amyloid A concentrations in antibiotic-treated calves.** Relationships between the relative abundance of faecal bacterial genera and serum amyloid A (SAA) concentrations were assessed in antibiotic-treated (ATB) calves using longitudinal correlation analyses. For each genus, the table reports the BRDC severity cluster in which the association was identified, the number of observations included in the analysis (Points), the estimated regression coefficient (Slope), the coefficient of determination (R²), and the Benjamini-Hochberg (BH)-adjusted P-value. Positive slope values indicate that higher genus abundance was associated with higher SAA concentrations, whereas negative values indicate an inverse relationship. Adjusted *P*-values account for multiple testing.

**Supplementary Table 6. Associations between nasal bacterial genera and serum amyloid A concentrations in non-antibiotic-treated calves** Associations between the relative abundance of nasal bacterial genera and serum amyloid A (SAA) concentrations were assessed in calves that did not receive antibiotic treatment (noATB). Analyses were performed separately within each BRDC severity cluster (Low, Moderate, and Severe). For each genus, the table reports the severity cluster in which the association was detected (Cluster), the number of observations included in the analysis (Points), the estimated regression coefficient (Slope), the coefficient of determination (R²), and the Benjamini-Hochberg (BH)-adjusted P-value. Positive slope values indicate that higher relative abundance of the genus was associated with higher SAA concentrations, whereas negative slope values indicate an inverse relationship. Adjusted *P*-values account for multiple testing correction using the BH procedure.

**Supplementary Table 7. Effects of antibiotic treatment and pathogen exposure on faecal short-chain fatty acid concentrations** Post hoc pairwise contrasts (estimated marginal means, emmeans) derived from linear mixed-effects models evaluating the effects of antibiotic treatment (ATB vs noATB) and pathogen exposure (Inf vs noInf) on faecal short-chain fatty acid (SCFA) concentrations. The table reports model estimates, standard errors (SE), degrees of freedom (df), test statistics (*t*-ratio), and associated *P*-values for acetate, propionate, butyrate, isobutyrate, valerate, and isovalerate. Estimates are presented on the model scale. Significant *P*-values indicate differences in SCFA concentrations between treatment or exposure groups. *P*-values < 0.05 were considered statistically significant

## Supplementary Figures

**Supplementary Figure 1.**
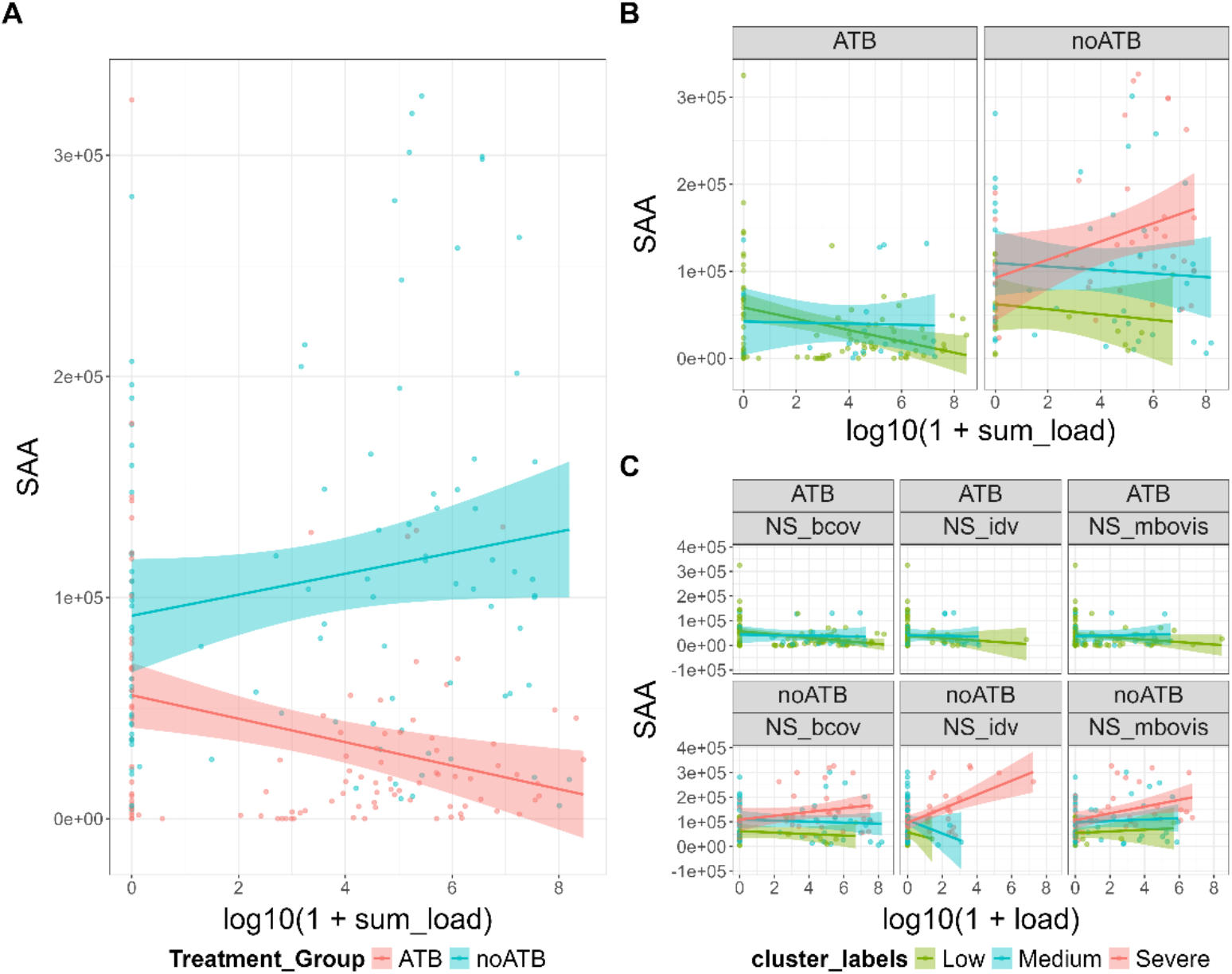
Serum Amyloid A blood concentration. **(A)** SAA blood concentration per total pathogen load in nasal cavities, quantified using targeted qPCR, and per antibiotic treatment condition. **(B)** SAA blood concentration per total pathogen load and per BRDC severity outcome. **(C)** SAA blood concentration per pathogen load and per BRDC severity outcome.

**Supplementary Figure 2.**
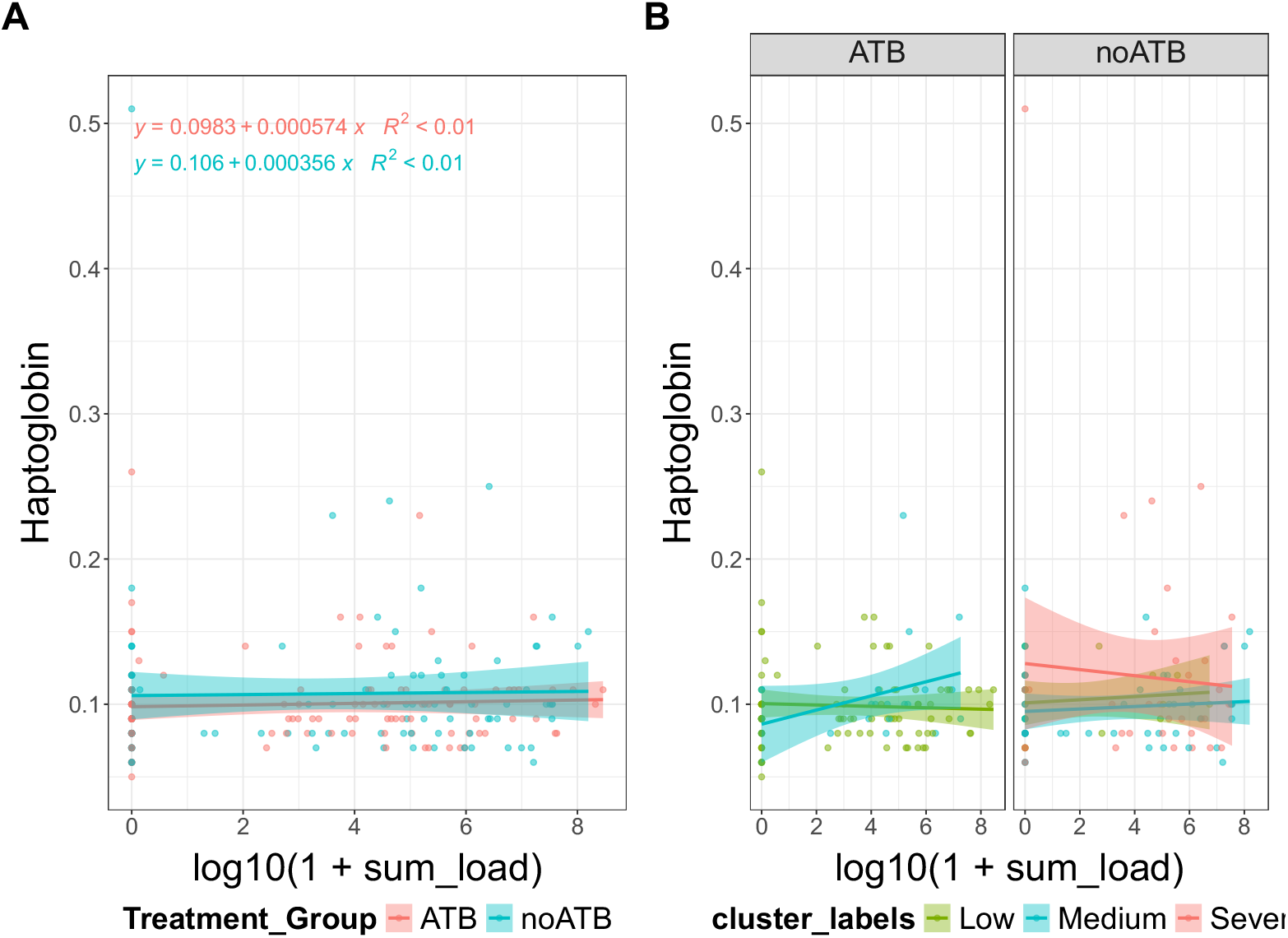
Haptoglobin blood concentration. **(A)** Haptoglobin blood concentration per total pathogen load in nasal cavities, quantified using targeted qPCR, and per antibiotic treatment condition. **(B)** Haptoglobin blood concentration per total pathogen load and per BRDC severity outcome. **(C)** Haptoglobin blood concentration per pathogen load and per BRDC severity outcome.

**Supplementary Figure 3.**
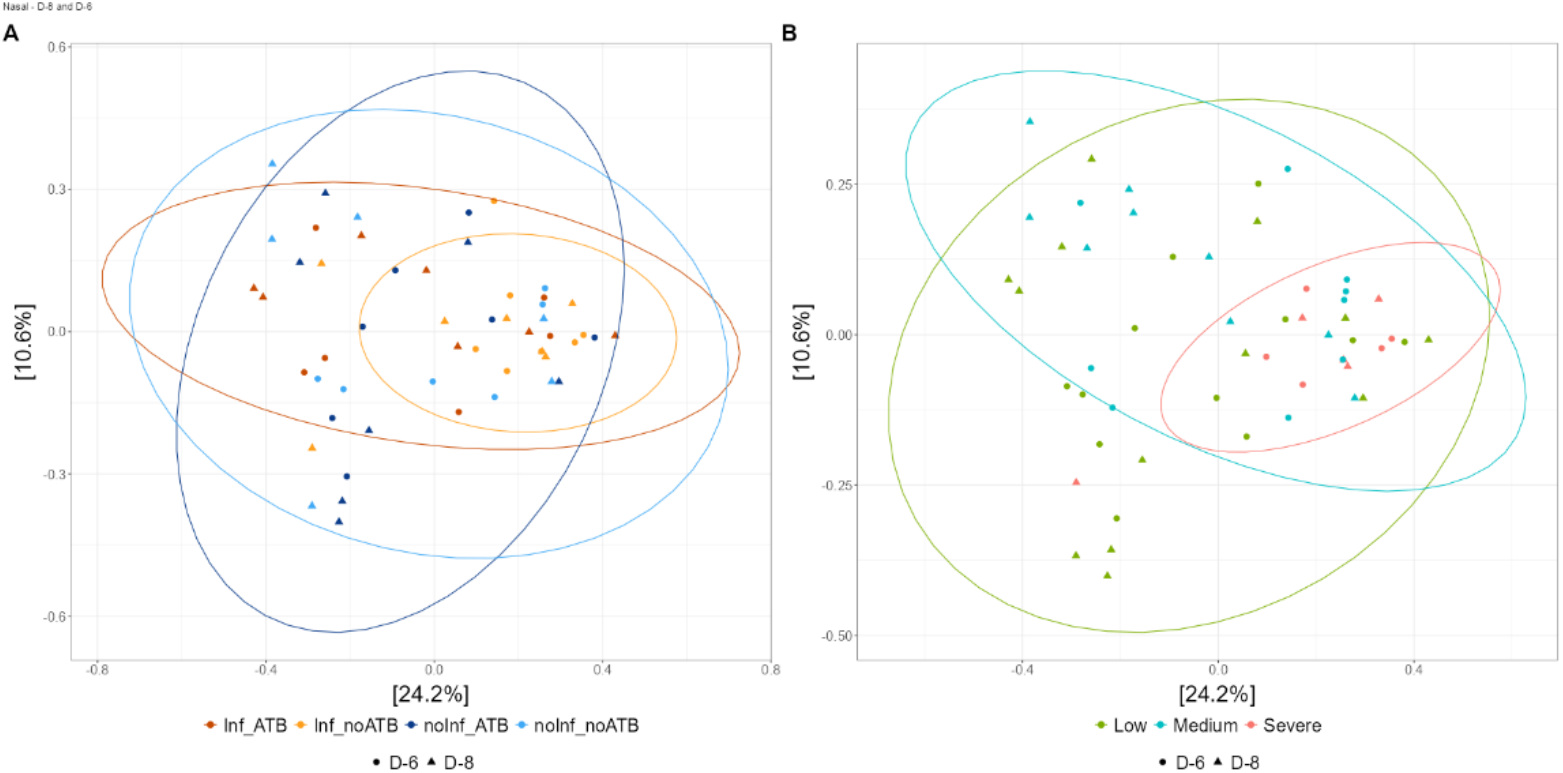
Nasal microbiota early-day Bray-Curtis beta diversity. **(A)** Nasal microbiota early-day beta diversity per experimental condition. **(B)** Nasal microbiota early-day beta diversity per BRDC outcome.

**Supplementary Figure 4.**
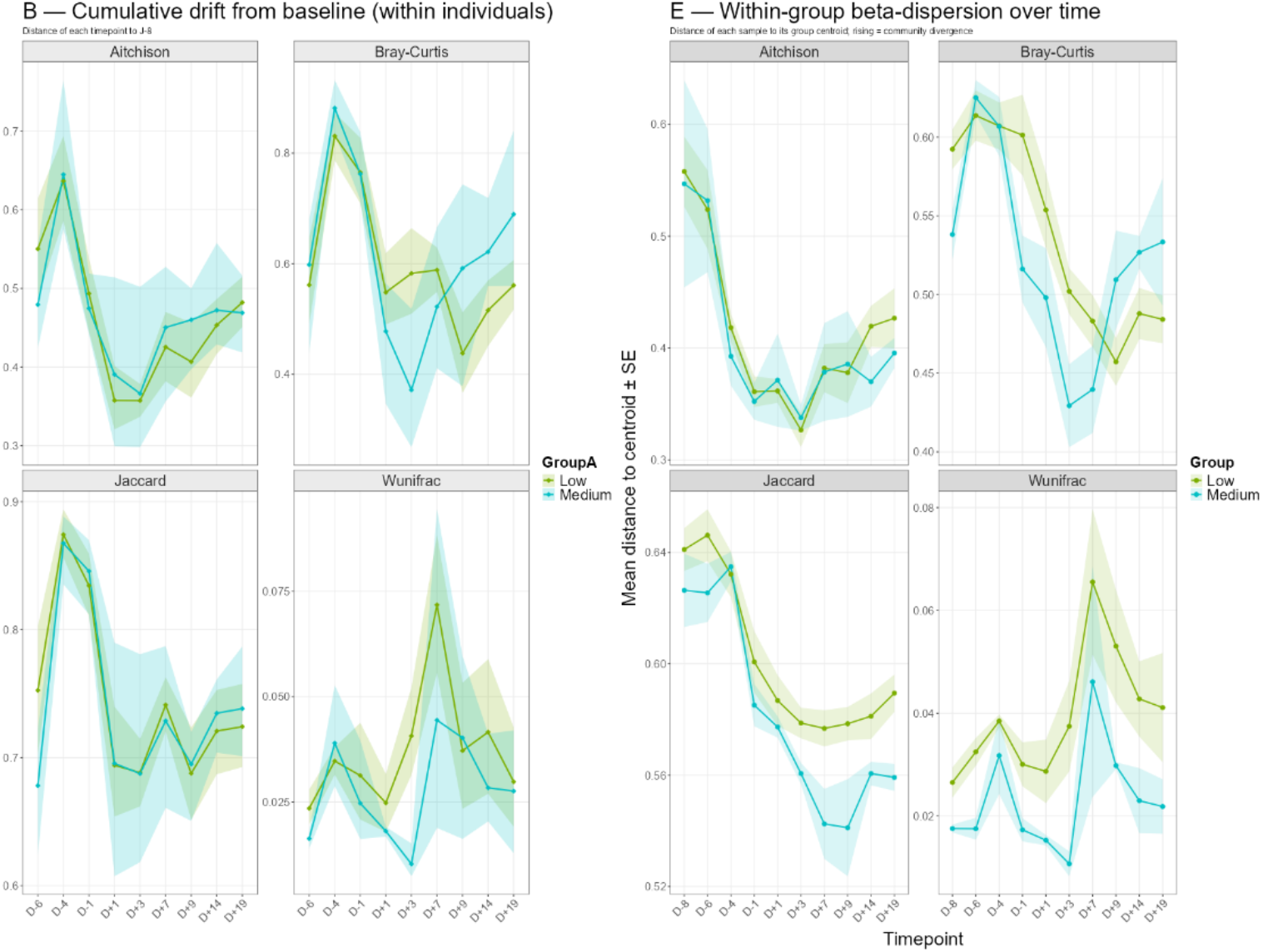
ATB-treated group faecal beta-diversity evolution. **(A)**. Beta-diversity: individual faecal microbiota distance from D-8. **(B)** Beta-diversity individual distance to the group centroid for each day.

**Supplementary Figure 5.**
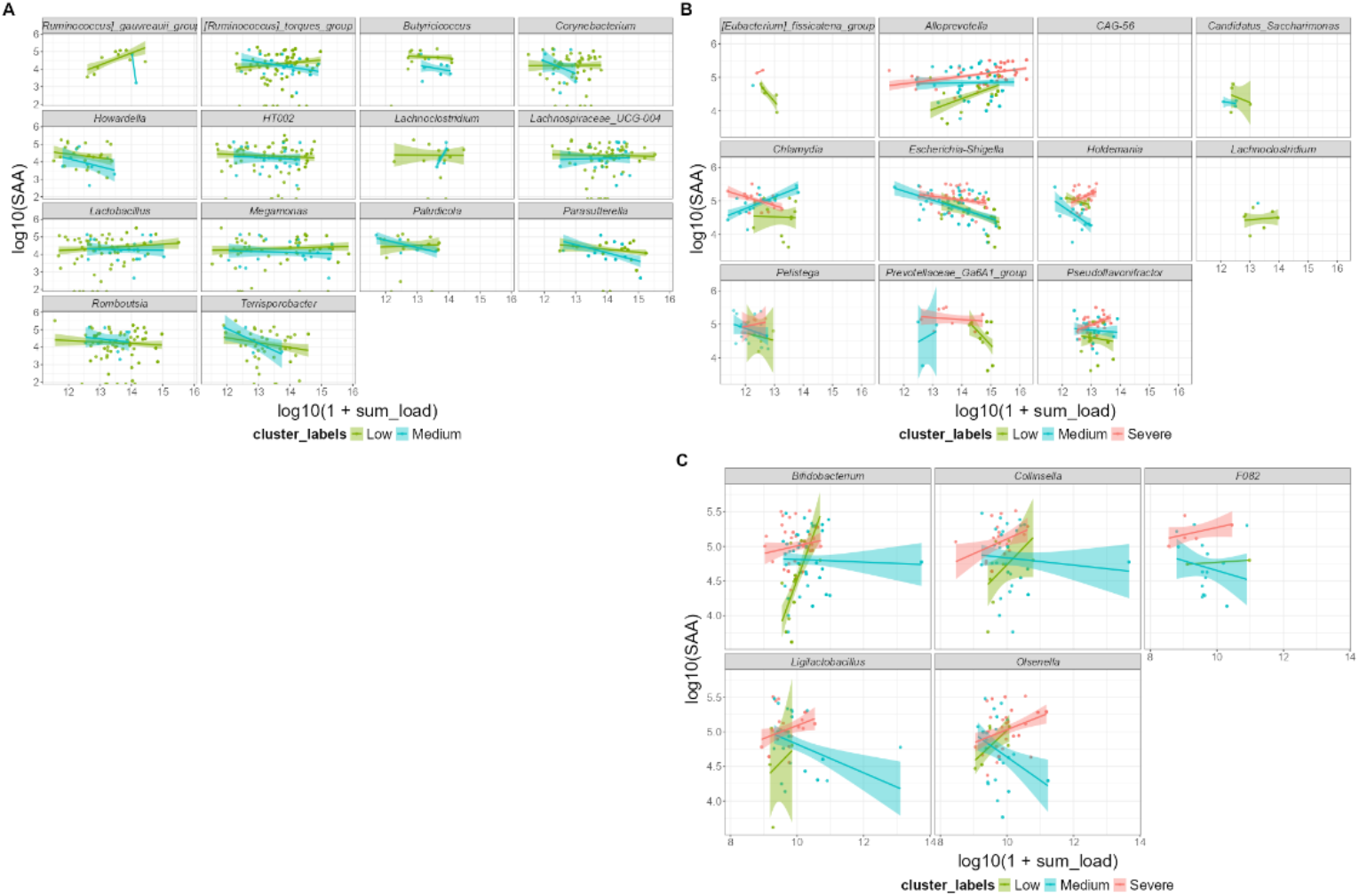
SAA and genera of interest. **(A)**. SAA concentration per genus of interest in the faecal microbiota of ATB-treated calves. **(B)** SAA concentration per genus of interest in the nasal microbiota of non-ATB-treated calves. **(C)** SAA concentration per genus of interest in the nasal microbiota of ATB-treated calves.

