## Supplementary figures and images for "Gut and nasal microbiome trajectories shape respiratory disease severity in calves exposed to antibiotic and pathogen challenge"

### Figure S1

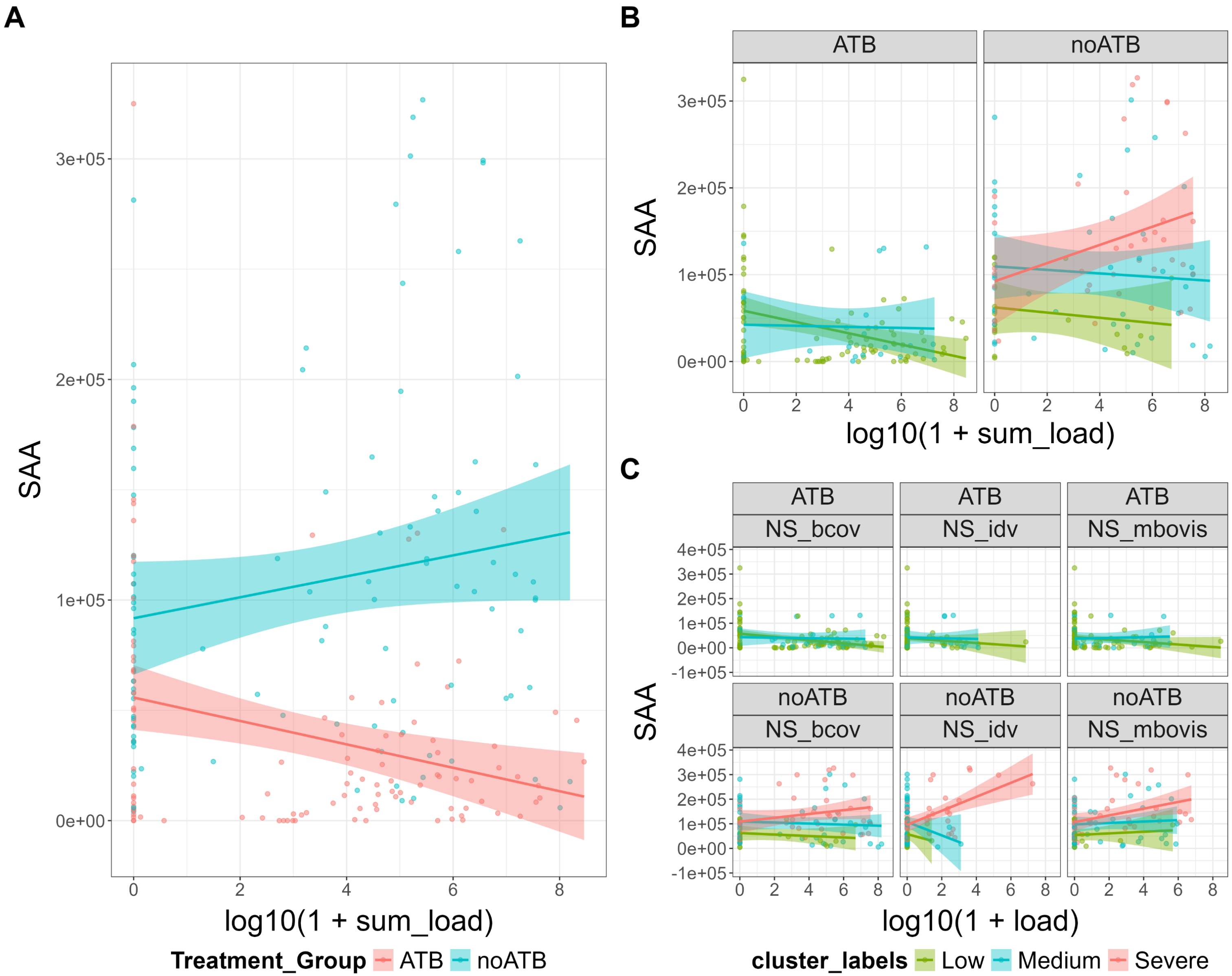

### Figure S2

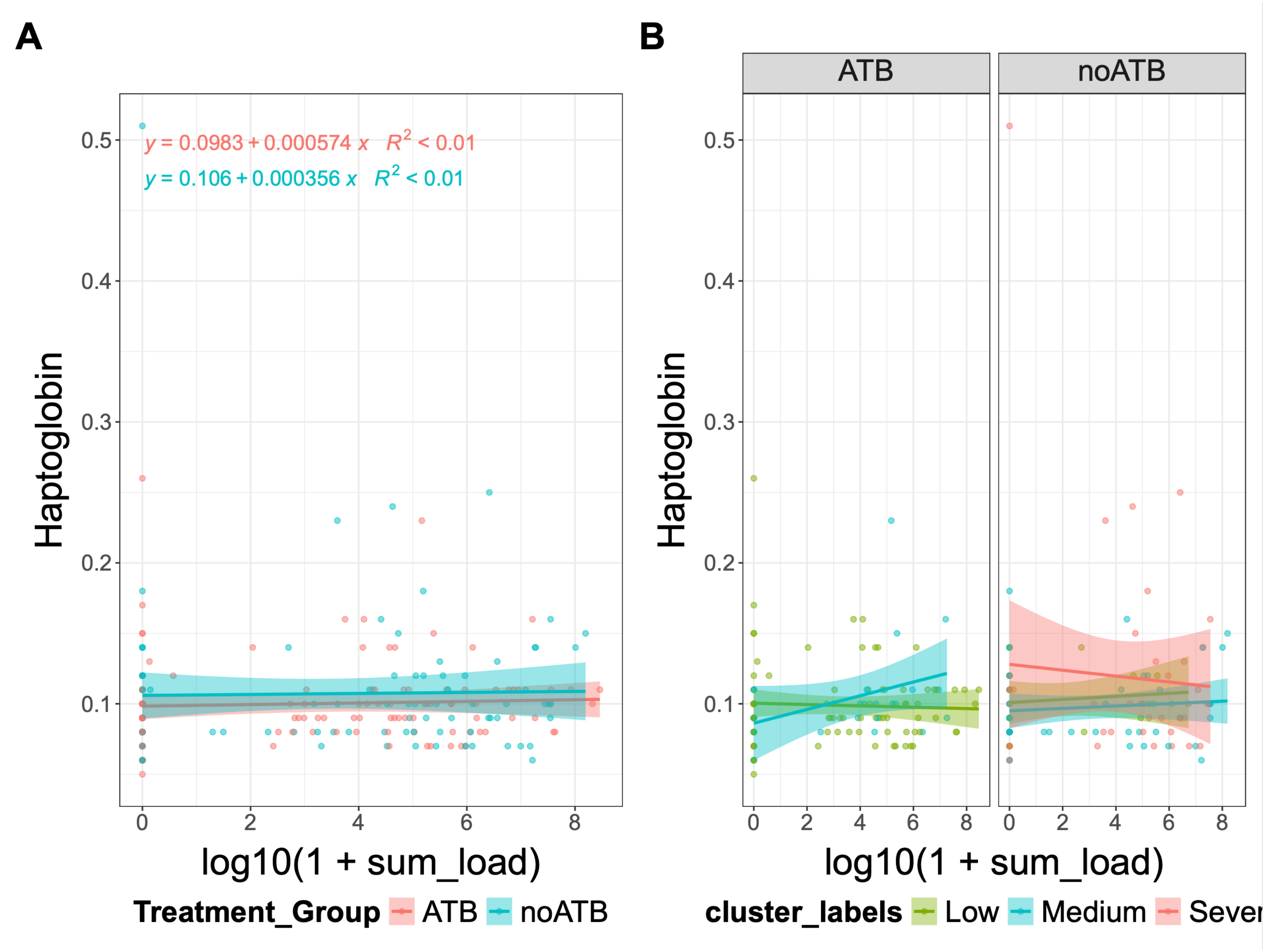

### Figure S4

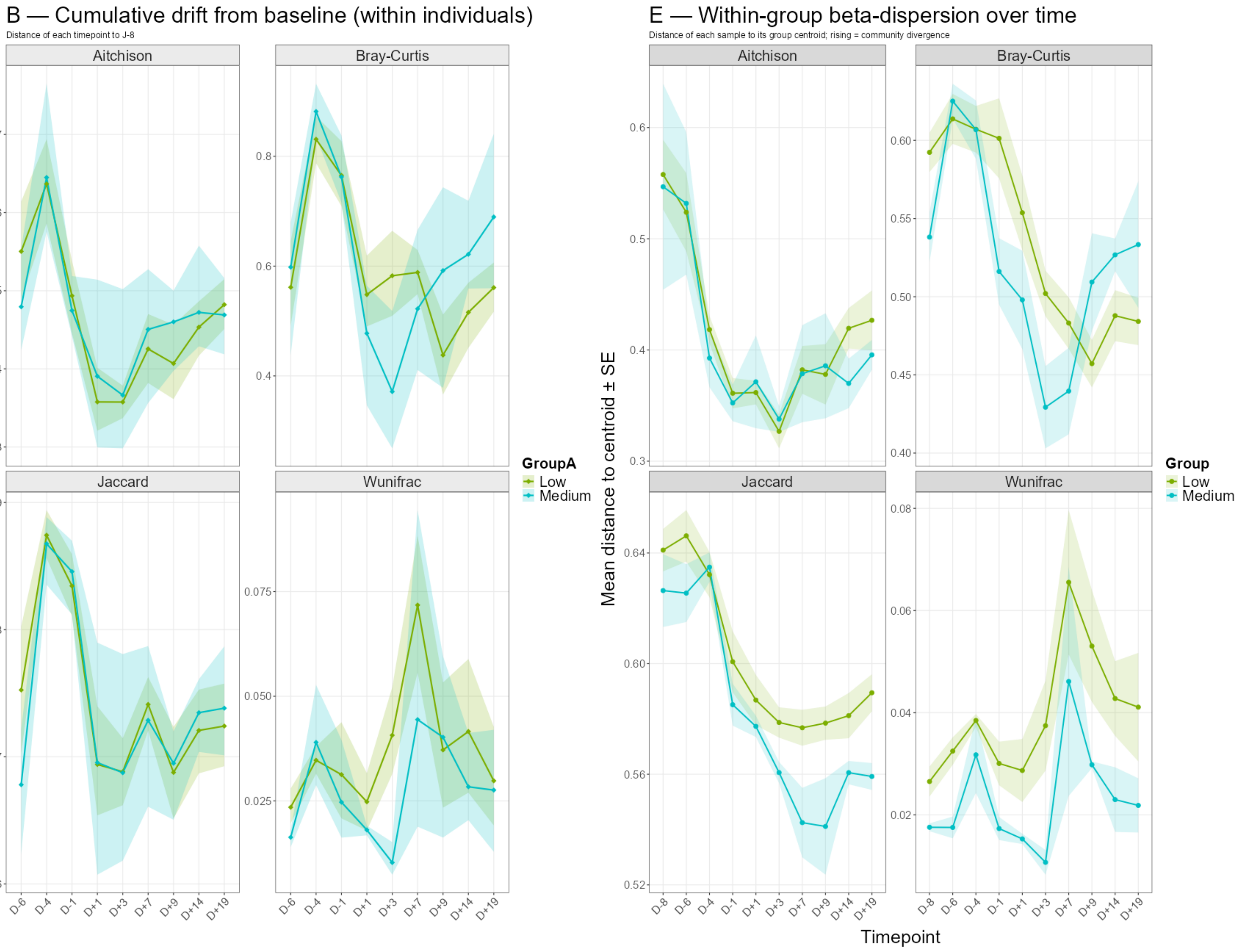

### Figure S5

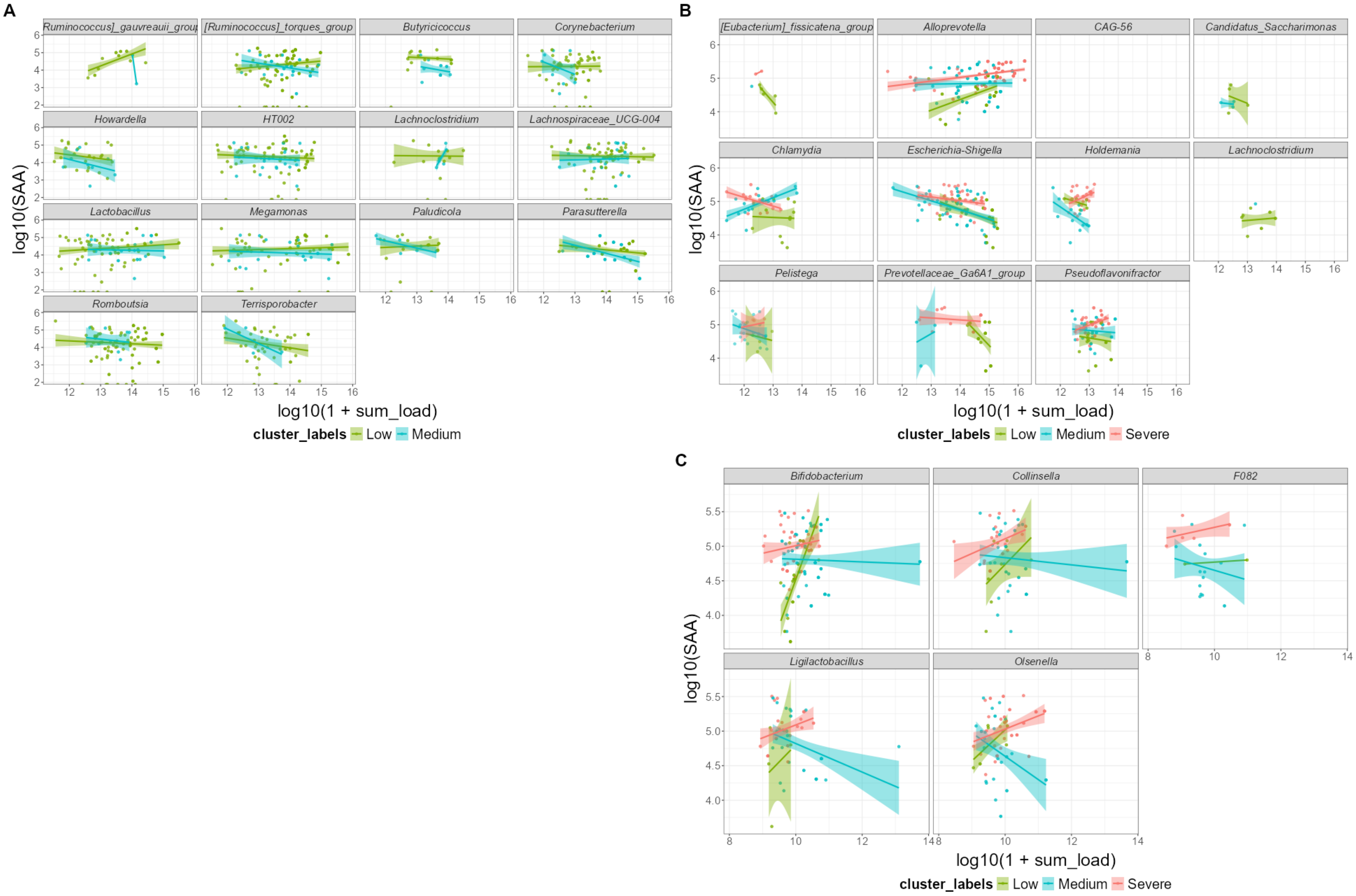
