## Supplementary material for "Gut and nasal microbiome trajectories shape respiratory disease severity in calves exposed to antibiotic and pathogen challenge": Figure S3

However, an analysis focused on the early-day microbiota (D-8 and D-6) revealed a difference in nasal microbiota composition homogeneity between BRDC severity

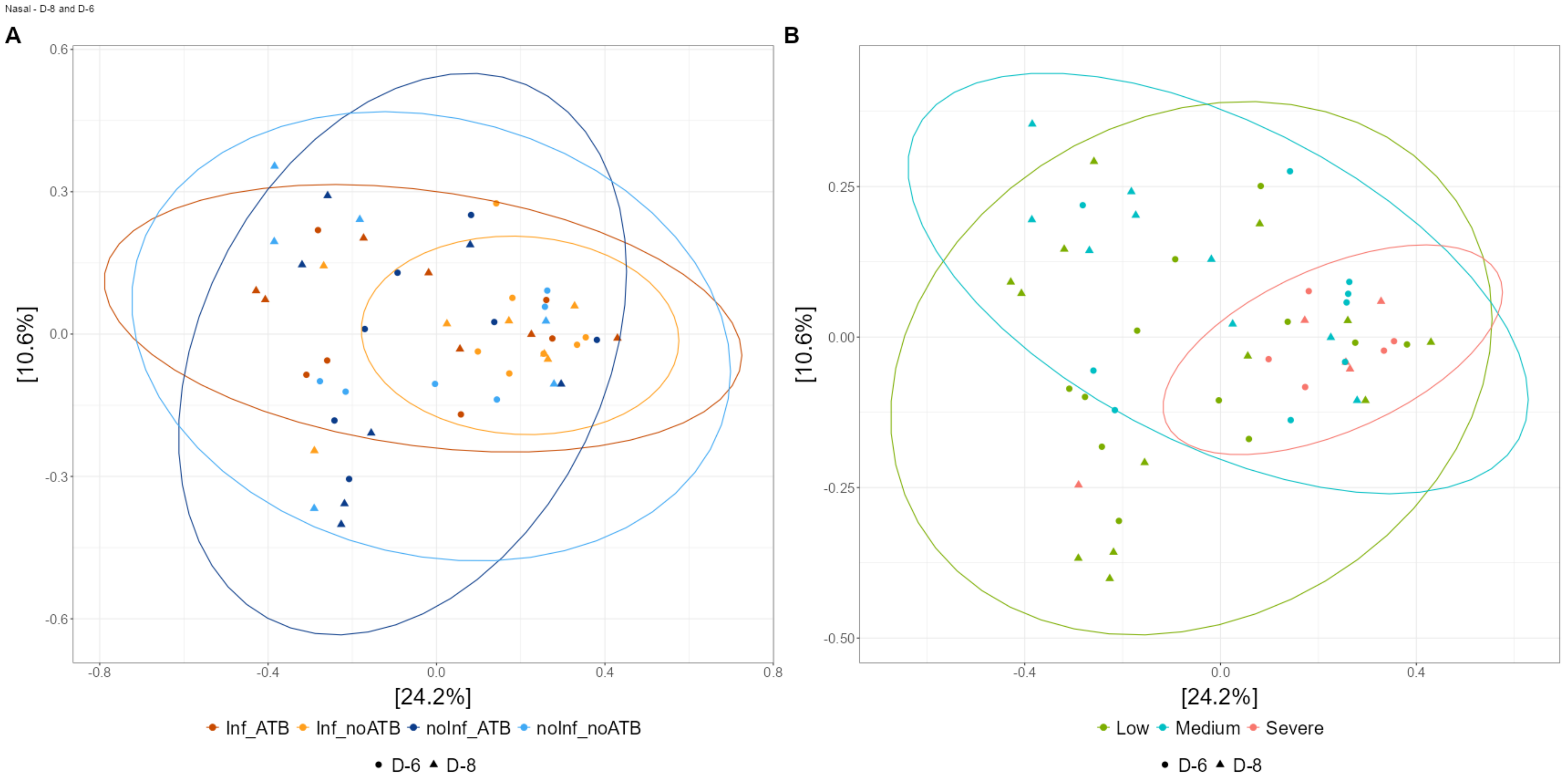
